# Data integration with uneven spatiotemporal coverage: A point-process approach for dynamic species distribution models

**DOI:** 10.1101/2025.11.07.687170

**Authors:** Moritz Klaassen, Marc Fernandez, Finn Lindgren, Virginia Morera-Pujol, Len Thomas, Nuno Oliveira, Joana Castro, Francisco Martinho, Sara Magalhães, Alfredo Rodrigues, Miguel P. Martins, Filipe Alves, Tiago A. Marques

## Abstract

Species distribution models (SDMs) are widely used to predict where and when species occur, but the data available to fit them often resolve space and time unevenly. Designed surveys, such as line-transect distance-sampling surveys, provide structured spatial coverage and the detection information needed to estimate detectability, but logistical and funding constraints often limit them to short, infrequent periods. Opportunistic or citizen-science records are collected more frequently across the year, but from areas shaped by observer access and interest rather than survey design, and usually without comparable detection information. As a result, models built from either source alone tend to capture spatial structure or temporal dynamics well, but not both, limiting predictions for mobile species in dynamic environments. Here, we present a general framework, based on a log-Gaussian Cox process (LGCP) fitted using integrated nested Laplace approximations (INLA), that links multiple data sources to a shared spatiotemporal measure of species density through source-specific observation models: a detection function for survey transects, and spatial constraints reflecting the footprint of opportunistic effort. We apply the framework to cetacean line-transect surveys and whale-watching records. In a simulation with one month of survey data and twelve months of opportunistic data, the integrated model recovered spatiotemporal distribution patterns well (median correlation with the true pattern = 0.91) and estimated monthly abundance with a mean error of +10.7%, outperforming single-source models. In a case study of common dolphins (*Delphinus delphis*) off mainland Portugal, integration improved the spatial accuracy of predictions, although weak covariate effects limited the temporal variation the model could resolve. These results show that integration adds the most value when the two data sources resolve complementary aspects of distribution, in this case broad spatial structure from the survey and temporal replication from whale-watching, while temporal predictions remain limited when covariates carry weak seasonal signal. The framework is general and can be extended to other taxa, systems, and combinations of structured and opportunistic data by substituting the relevant observation models.

## 1 Introduction

Spatial models of abundance and distribution, often collectively referred to as species distribution models (SDMs), are a core tool for predicting where, and increasingly when, species occur (Elith and Leathwick 2009). They are applied across terrestrial, marine, and freshwater systems, and the predictions they produce increasingly inform conservation and management decisions that depend on how distributions shift through time, such as the dynamic management of mobile species in changing environments (Hazen et al. 2018). These models can be fitted to data from a wide range of sampling methods and observation modalities, including line-transect distance sampling, point counts, camera traps, telemetry, environmental DNA, and citizen-science records (Miller et al. 2019). The increasing availability and variety of these data have, in turn, motivated methods that combine more than one data source within a single modelling framework — see, e.g., Pacifici et al. (2017), which was featured in the recent virtual collection “A Century of Statistical Ecology” (Gilbert et al. 2024).

Designed surveys follow a sampling protocol that connects what is observed to the underlying population, typically by recording effort and the information needed to model imperfect detection, such as the distances at which animals are detected (Buckland et al. 2001). Opportunistic records, such as citizen-science reports or sightings logged by ecotourism operators, instead arise as a byproduct of activity that no survey design governs. As a result, the spatial distribution of effort, its intensity, and the detectability of animals are generally unknown and cannot be modelled directly from the data alone (Renner and Warton 2013). Because designed surveys require dedicated effort, logistical and funding constraints frequently restrict them in space, in time, or both. Opportunistic records can accumulate over wide areas or long periods, but concentrate wherever observers happen to go. Whether a survey is broader in space than in time, or the reverse, and whether opportunistic records compensate in space or in time, depends on the particular survey and platform rather than on the data type itself. What determines whether combining them is worthwhile is whether the two sources are complementary: whether one resolves spatial structure the other lacks, while the other resolves temporal change the first cannot.

A concrete and instructive instance motivates the framework we present. Many mobile marine species, including cetaceans, are monitored by shipboard and aerial line-transect distance sampling, the most widely used method for assessing cetacean occurrence (Kaschner et al. 2012). These surveys record where groups of animals are seen, along with the detection distances needed to estimate their abundance. Such surveys are expensive and logistically demanding, so they are typically confined to short summer windows when sea state and daylight allow reliable detection. The SCANS programme illustrates this: these surveys cover large areas of the European Atlantic continental shelf, yet each is completed within a single summer, and the full programme has been carried out only four times in nearly three decades (Hammond et al. 2002; Hammond et al. 2013). Each survey therefore yields a detailed spatial snapshot and a robust abundance estimate, but reveals little about how distributions change between the surveys. The same constraint applies at regional scales: surveys of this kind are typically confined to a single season regardless of their spatial extent, so the temporal gap arises at the finer scales at which monitoring and management most often operate. The same species targeted by these surveys are often recorded throughout the year by opportunistic sources: whale-watching operators, vessels of opportunity, and passive acoustic monitoring programmes, among others (Klaassen et al. 2025). These sources can accumulate many observations across time at comparatively low cost, but each reflects a distinct observation process, shaped by its own spatial footprint, effort distribution, and detection characteristics.

Whale-watching operators, for example, report sightings from the coastal areas they routinely visit but provide no information on detectability or on the distribution of animals beyond those areas. Neither source alone can describe how the distribution and abundance of animal groups change from month to month across the wider region: the survey spans the region but only for a single month, while the whale-watching records span the year but only along the coast.

To integrate these complementary but unevenly resolved data sources, we develop a spatiotemporal SDM that couples a distance-sampling likelihood with a presence-only likelihood through a shared latent intensity surface, modelled as a log-Gaussian Cox process (LGCP) and fitted with integrated nested Laplace approximations (INLA). Such joint-likelihood frameworks are now well established (Kéry et al. 2024; Farr et al. 2021; Martino et al. 2021), and spatiotemporal approaches are emerging as well (Seaton et al. 2024; Farr et al. 2024). Here, we address a setting that to our knowledge has not been explored: a structured survey covering the full region but conducted only once, paired with opportunistic records that span a much longer period but only a small fraction of that region, with the aim of estimating group density across the full spatiotemporal domain. The distance-sampling component here models detectability explicitly, so the survey anchors absolute group density. Rather than estimating a residual spatial field that itself varies through time (Seaton et al. 2024), we retain a single shared spatial field and carry temporal variation through dynamic covariates, a structure suited to settings where the broad structured survey covers only a single period.

We evaluate the framework through a simulation study that asks not only whether the integrated model recovers known spatiotemporal structure, but when it adds value over either source alone. We then apply it to a case study of common dolphins (*Delphinus delphis*) off mainland Portugal. The framework is general: any combination of a structured survey with a tractable detection model and a temporally richer opportunistic source can be accommodated by substituting the relevant observation models.

## 2 Material and Methods

### 2.1 Modelling approach

We model the distribution of animal groups as a log-Gaussian Cox process (LGCP): groups are distributed in space and time as an inhomogeneous Poisson point process whose log-intensity is the sum of a linear predictor in environmental covariates and a Gaussian random field (GRF) capturing residual spatial structure (Illian et al. 2008). Each dataset observes this process through its own observation model, shaped by its spatial coverage, sampling protocol, and detection characteristics. The joint model estimates the shared intensity while accounting for these differences through dataset-specific likelihoods, allowing covariate responses and residual spatial structure to be shared across datasets while scale differences and detectability are handled separately. In this application, the model operates at monthly temporal resolution, combining one month of structured survey data with twelve months of opportunistic records.

#### 2.1.1 Latent spatiotemporal density and absolute abundance

Let **s** ∈ Ω ⊂ ℝ^2^ denote a two-dimensional spatial location, and let *t* ∈ *T* = {1, … , 12} index months (January to December). The latent density *λ*(**s**, *t*), representing the expected number of animal groups per unit area at location **s** in month *t*, is

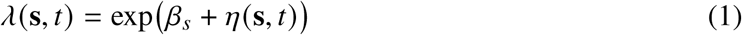

where *β_s_* is the survey intercept that places *λ*(**s**, *t*) on an absolute group-density scale and *η*(**s**, *t*) is the shared linear predictor. By integrating the latent density *λ*(**s**, *t*)) over the entire spatial domain Ω, we obtain the absolute abundance of animal groups N(t) for a given month

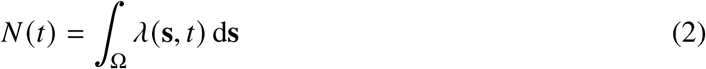

We write the linear predictor as

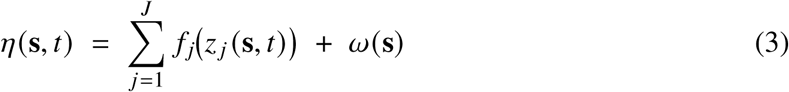

where *J* is the number of covariates, *z _j_* is the value of the *j*th environmental covariate, *f _j_* is its smooth response, and *ω*(**s**) is a Matérn Gaussian random field implemented via the SPDE approach. We use a purely spatial residual field, *ω*(**s**), rather than a spatiotemporal field *ω*(**s**, *t*). Temporal variation is represented by the time-varying covariates *z _j_* (**s**, *t*). This reflects the present data configuration and is discussed further in §4.1.

#### 2.1.2 Spatial field

We represent *ω*(**s**) as a continuous-space Matérn GRF, formulated through the stochastic partial differential equation (SPDE) representation (Lindgren et al. 2011). This formulation enables a sparse Gaussian Markov random field (GMRF) approximation on a triangulated mesh, providing computational efficiency while retaining the properties of the underlying GRF. The Matérn covariance between two locations separated by distance r = ‖s − s′‖ is

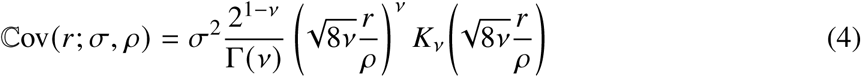

where *K_ν_* is the modified Bessel function of the second kind, *σ* is the marginal standard deviation, *ρ* is the practical range (Euclidean distance where correlation is ≈ 0.1), and *ν* is the smoothness parameter, which is fixed to 1. We used penalised complexity (PC) priors on ( *ρ*, *σ*), specified by tail probabilities ℙ( *ρ* < *ρ*_0_) = *α*_1_ and ℙ(*σ* > *σ*_0_) = *α*_2_ (Simpson et al. 2017). Where appropriate, we employ a barrier variant of the SPDE to prevent correlation across land masses (Bakka et al. 2019). Covariate responses *f _j_* (·) are modelled as nonlinear 1-D SPDE smooths with PC priors, using the same Matérn representation as the spatial field; details are given in Appendix S1: Section S2.

#### 2.1.3 Observation models

We link the different data sources to the shared latent group density *λ*(**s**, *t*) via their observation-specific point-process likelihoods.

##### (i) Line-transect survey with distance sampling

We formulate our model building on the conventional distance-sampling observation process (Buckland et al. 2015) where detections of animal groups along transect lines are modelled as an inhomogeneous Poisson point process thinned by a detection function. Let *t* be the survey month (fixed for this likelihood), *C* ⊂ Ω represent the total area covered by the survey strips in that month (defined by a truncation half-width *W*), and *d*(**s**) the perpendicular distance from **s** to the nearest transect line. The line-transect log-likelihood is specified as

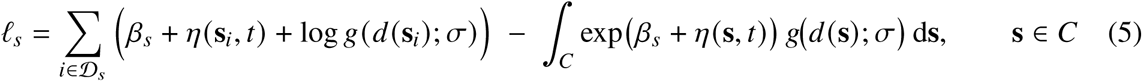

where *β_s_* is the survey-specific intercept, *g*(·) is the detection function, and *D_s_* indexes observed animal groups. We adopt the half-normal key for the detection function as the main specification,

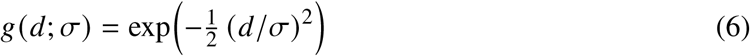

with a weakly informative prior distribution on the scale parameter *σ* set via an exponential mapping on the natural scale (centred relative to *W*). As a sensitivity check for the case study, we also fitted a hazard-rate detection function with the shape parameter fixed to *b* = 1:

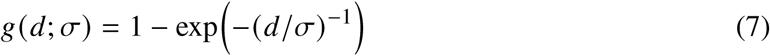

##### (ii) Whale watching

We treat opportunistic whale-watching (WW) sightings of animal groups as a presence-only point process restricted to an active commercial search domain. We assume the defining feature of this data source to be concentrated, highly coordinated searching effort: multiple observers scan simultaneously (from vessels and shore lookouts), operators communicate by radio, follow each other to sightings, and commercial incentives (e.g., refund policies) favour thorough search within operating footprints. Rather than attempting to reconstruct a fine-scale effort surface for many unknown, overlapping, and mobile platforms we condition the likelihood on spatial availability. We define this availability as the geographic footprint where commercial whale-watching operations actually took place. In practice, for each month *t* ∈ {1, … , 12}, we construct an availability domain *A*(*t*) ⊂ Ω as the convex hull of boat GPS positions recorded in that month, or of sightings where GPS tracks were unavailable. These domains vary by month, reflecting the seasonal pattern of whale-watching operations: footprints usually expand during the summer peak and contract in winter. Restricting the likelihood integral to *A*(*t*) prevents unvisited areas from being treated as surveyed with zero detections: because whale-watching operators did not cover the full spatial domain Ω, the absence of sightings outside *A*(*t*) carries no information about group density there. Month-to-month variation in total search effort is captured through a known scalar *q_t_* = *n*_op_(*t*), the number of days on which whale-watching boats operated in month *t*. This quantity is available from operational records and enters the likelihood as a fixed offset, accounting for the fact that vessels operate more frequently during favourable summer conditions than in other months. The WW likelihood is then given by

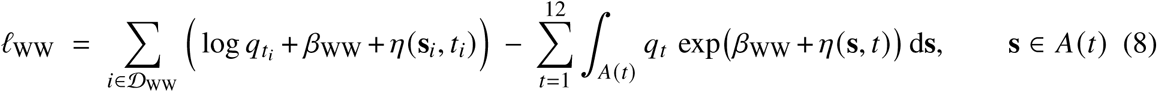

where *q_t_* > 0 is the month-specific exposure scalar (*q_t_* = *n*_op_(*t*), the number of operating days in month *t*), *β*_WW_ is the WW intercept, *D*_WW_ indexes observed animal groups from WW, and *A*(*t*) are month-specific availability polygons.

##### (iii) Joint log-likelihood and posterior

Let *y* = (*y_s_*, *y*_WW_) be the observed data, 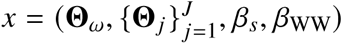 the latent Gaussian variables (**Θ***_ω_* are SPDE weights for *ω*(**s**); **Θ***_j_* are 1-D SPDE weights for *f _j_* (·)), and 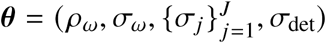 the hyperparameters, where ( *ρ_ω_*, *σ_ω_*) are the Matérn range and marginal SD of *ω*(**s**), *σ_j_* are the marginal SDs of the 1-D smooths, and *σ*_det_ is the detection-scale parameter in *g*. Assuming conditional independence given (*x*, ***θ***), the joint log-likelihood factorises as

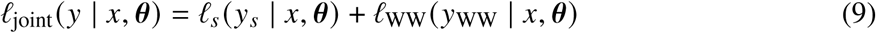

The log-posterior probability density of (*x*, ***θ***) given *y* is

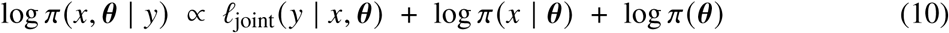

Uncertainty propagation through the joint posterior and model diagnostic checks are described in Appendix S1: Section S1.

#### 2.1.4 Inference and implementation

All analyses were run in R (version 4.4.2; R Core Team, 2024) with inlabru (Bachl et al. 2019), which builds on INLA (Rue et al. 2009) to fit latent Gaussian spatial models. Spatial (2-D), distance and covariate (1-D) meshes were built with fmesher (Lindgren 2025), and barrier SPDEs used INLAspacetime (Krainski et al. 2025). A full list of packages used, and their versions, can be found in Appendix S1: Table S2.

### 2.2 Simulation study

We used a simulation study to assess whether the joint model recovers known space–time structure under a specific data configuration (Figure 1). For that, we generated the distribution of a virtual species for the Azores (central North Atlantic) using non-linear responses to three environmental covariates: bathymetry, seafloor slope, and monthly SST, combined with a GRF, producing monthly suitability surfaces and targeting a group density of 0.4 groups per 100 km^2^. Survey detections were simulated for August only (*n* = 79) after half-normal thinning. WW availability polygons were informed by observed effort patterns in the region. The spatial field was discretised on a barrier-SPDE triangulated mesh with 8 km hexagonal seeding over the inner domain. Full details of occurrence generation, observation sampling, and mesh construction are given in Appendix S1: Section S3.

**Figure 1.**
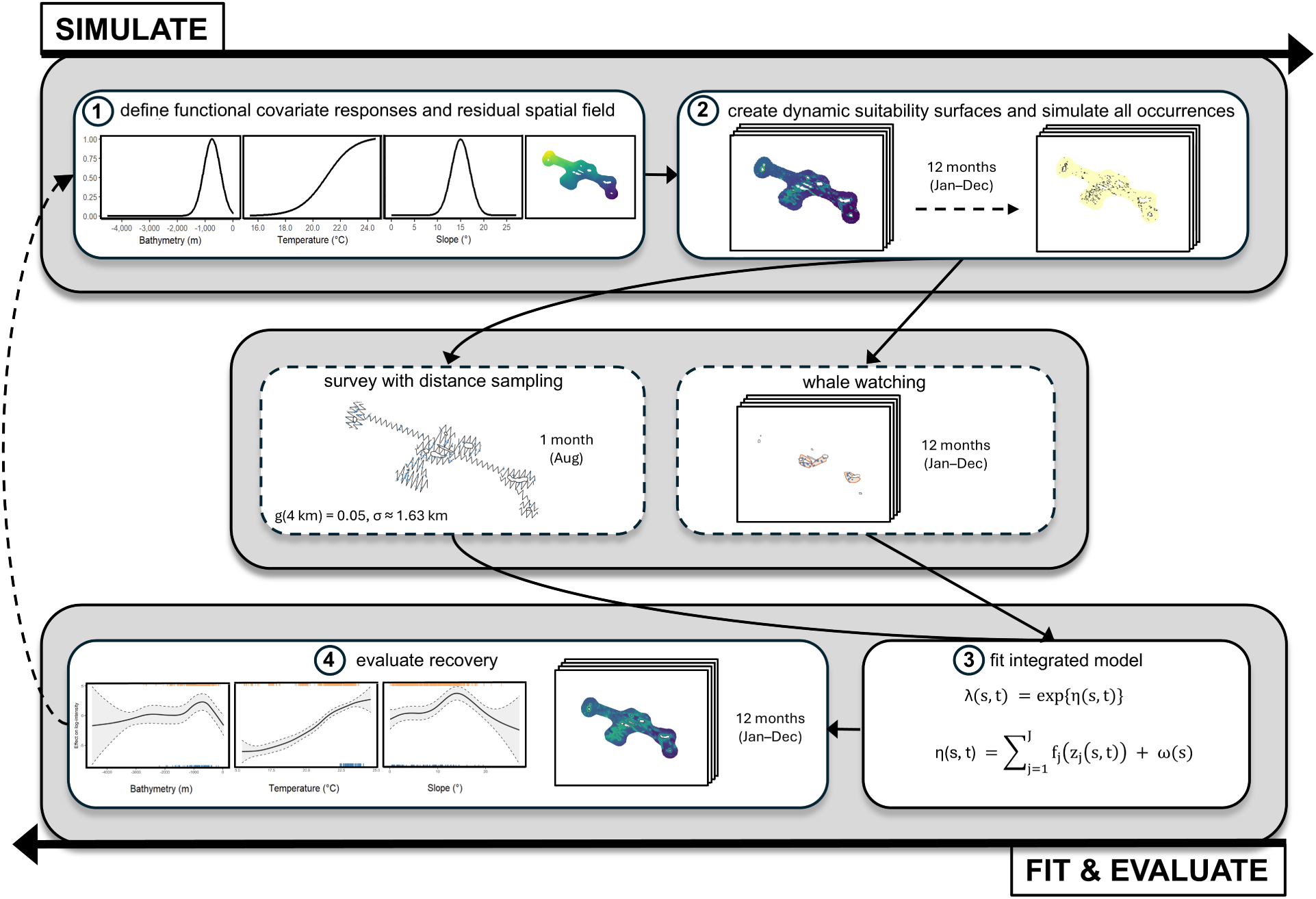
Design of the simulation. (1) Define functional covariate responses (bathymetry, temperature, slope) and a residual spatial field to generate (2) monthly habitat-suitability surfaces. Convert suitability to global occurrences for each month (n = 12, Jan–Dec). Sample the latent pattern through two observation processes (dashed): survey with distance sampling (August only) and whale-watching (WW) data restricted by monthly availability polygons (Jan-Dec). (3) Fit an integrated model with dataset-specific likelihoods. (4) Evaluate recovery of covariate effects, spatial field, monthly density maps, and absolute abundances.

**Table 1:** Monthly abundance (groups): simulation vs. model estimates.

| Month | Simulated mean (95% CI) | Model mean (95% CrI) | Relative error (%) |
| --- | --- | --- | --- |
| Jan | 33.1 (31.3, 34.9) | 38.5 (23.3, 60.1) | 16.4% |
| Feb | 32.4 (30.6, 34.1) | 37.1 (21.5, 55.9) | 14.5% |
| Mar | 32.2 (30.4, 34.0) | 34.6 (20.1, 55.2) | 7.4% |
| Apr | 32.2 (30.4, 33.9) | 35.1 (20.8, 54.7) | 9.0% |
| May | 34.7 (32.8, 36.5) | 41.6 (25.0, 65.0) | 20.0% |
| Jun | 60.2 (58.5, 62.0) | 74.1 (48.3, 107.8) | 23.0% |
| Jul | 232.9 (228.1, 237.8) | 226.5 (159.8, 309.7) | −2.7% |
| Aug | 305.8 (301.0, 310.5) | 284.9 (211.2, 375.4) | −6.8% |
| Sep | 302.2 (297.3, 307.1) | 282.1 (206.7, 371.5) | −6.7% |
| Oct | 148.1 (144.4, 151.8) | 161.1 (112.2, 227.6) | 8.7% |
| Nov | 66.5 (64.3, 68.7) | 80.0 (53.6, 116.7) | 20.3% |
| Dec | 37.9 (35.9, 39.8) | 47.4 (29.3, 73.6) | 25.0% |

*Notes:* Simulation values are 30-day means with 95% confidence intervals (CIs). Model values are posterior means with 95% credible intervals (CrIs). To smooth out Monte Carlo error, we used 2,000 posterior draws per month. The resulting numerical error was <5% of the SD. Relative error (%) was computed as $(\text{model} - \text{simulated}) / \text{simulated} \times 100$ .

#### 2.2.1 Quantifying the gain of data integration

The baseline simulation illustrates the joint-likelihood workflow under a single configuration. To test robustness and to quantify when data integration adds value, we extended the simulation to a grid of scenarios spanning variation in latent spatial structure and in the extent to which the ecological process is explained by covariates. These two axes capture the key sources of variation in how much information the two data sources share: the spatial scale at which unexplained structure varies, and the degree to which distribution is driven by measured covariates versus residual spatial pattern. We considered nine scenarios defined by two factors: (i) the correlation range of the GRF, *ρ*_water_ ∈ {100, 300, 600} km, and (ii) the relative contribution of measured covariates versus the residual field (patterns C3S1, C2S2, C1S3). Shorter *ρ*_water_ yields more locally varying residual structure, whereas longer *ρ*_water_ produces smoother broad-scale gradients. The covariate-field patterns were set so that covariates contribute three quarters of the linear predictor to the point-generating surface (C3S1), both contribute equally (C2S2), or the residual field contributes three quarters (C1S3). For each scenario we generated *n* = 30 independent replicates of the full simulation pipeline. Because the simulation outcomes depend on the GRF, and this field is itself a random realisation, we use these replicates to average performance over GRF variability rather than relying on a single draw (Appendix S1: Figure S10). For every simulated dataset we fitted three models: (i) the integrated model combining survey and WW likelihoods (Equation (9)), (ii) a WW-only model fitting only the WW likelihood (Equation (8)), and (iii) a survey-only model fitting only the survey likelihood (Equation (5)). This isolates the contribution of each data source to recovery of the underlying distribution. For each scenario, replicate, and month we computed group density surfaces on the analysis grid and evaluated recovery using Spearman’s rank correlation between the simulated suitability surface and the posterior mean log group density surface. Overall, the design yields 3 × 3 = 9 scenarios with *n* = 30 replicates and 12 monthly surfaces per replicate, producing 3240 month-specific correlations per model.

### 2.3 Case study

#### 2.3.1 Data

We applied the framework to (*D. delphis*) off mainland Portugal for 2020 at a monthly resolution. For the survey, we used a line-transect distance sampling survey in February from the region (Sociedade Portuguesa para o Estudo das Aves, SPEA) yielding *n* = 30 detections. The opportunistic component comprised WW data from four operational areas where commercial activities are run regularly: Lisbon (*n* = 54; Jan-Mar, Jun-Sep), Albufeira (*n* = 136; Jul-Nov, includes WW trips and non-systematic surveys, which we pooled in the analysis), Sagres (*n* = 269; Jan-Mar, Jun-Nov), and Faro (*n* = 83; Jan-Mar, Jun-Dec). For the Lisbon, Albufeira and Sagres datasets, time-ordered GPS fixes were used to build monthly availability polygons *A*(*t*). For Faro, monthly *A*(*t*) were constructed from sightings. Monthly sampling effort across all four operators is shown in Appendix S1: Figure S7.

#### 2.3.2 Modelling

We modelled the case study with a similar shared latent LGCP as above. To reflect *D. delphis* ecology, we used monthly means of SST and chlorophyll-*a* as dynamic predictors and slope as a static predictor. The survey likelihood used a half-normal detection function with truncation *W* = 0.25 km, implemented via the LGCP distance-sampling formulation (Equation (5)). As a sensitivity analysis, we also fitted a hazard-rate detection function; goodness-of-fit via posterior-predictive binned counts favoured the half-normal (Appendix S1: Figure S8).

## 3 Results

### 3.1 Simulated data

#### (i) Spatial patterns

Posterior mean log-density surfaces from the integrated model matched the simulation well (across months, median Spearman’s *ρ* = 0.91; IQR 0.89-0.91). The barrier SPDE was recovered (Figure 2A; cf. Appendix S1: Figure S1), and the month-to-month patterns followed the simulated temperature-driven seasonality from the point-generating suitability surface (Figure 2B; cf. Appendix S1: Figure S11).

**Figure 2.**
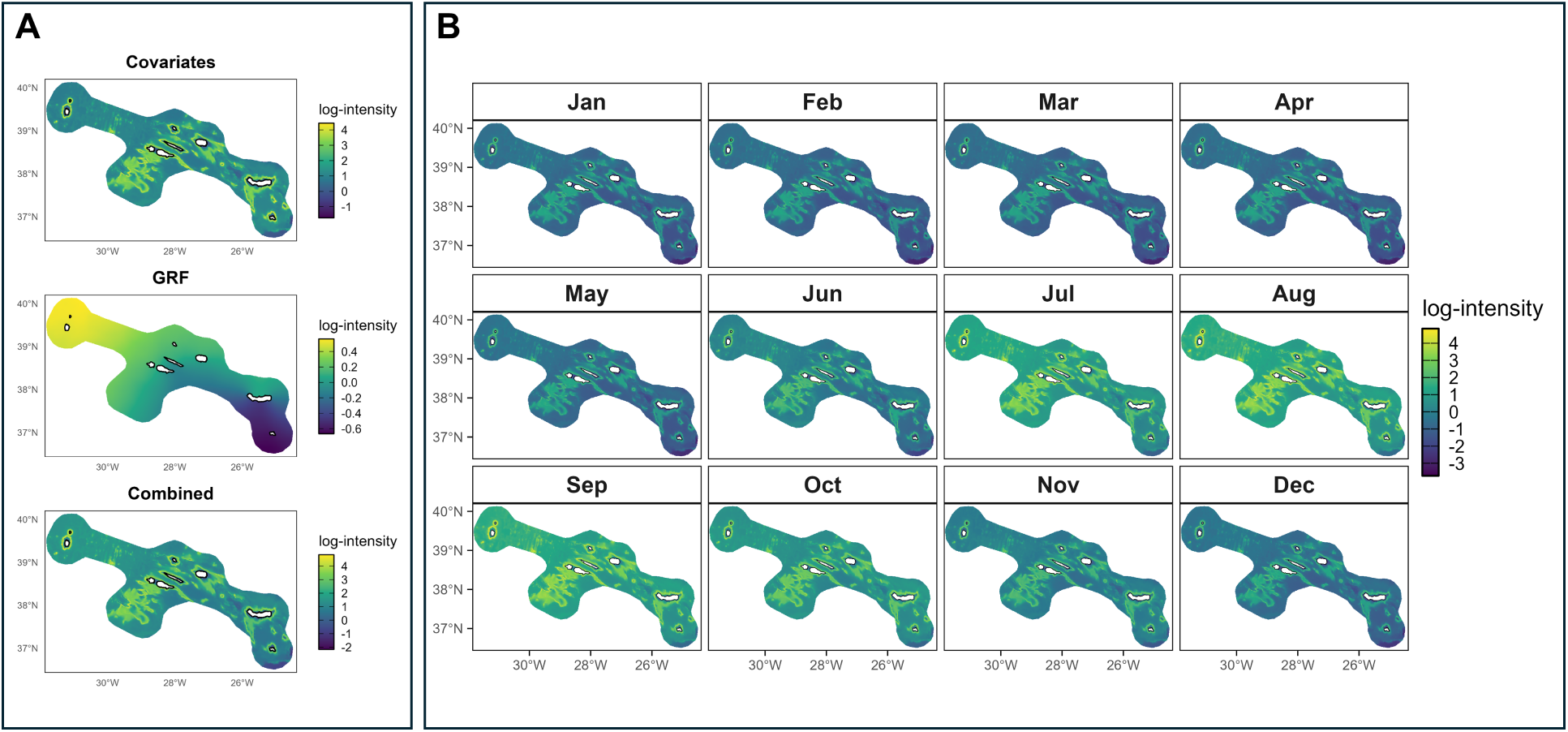
Posterior mean log-density estimated by the integrated model. **(A)** Model components for August (example month): covariates, spatial Gaussian random field (GRF), and the combined linear predictor. **(B)** Monthly log-density surfaces (Jan–Dec).

#### (ii) Component recovery

The joint-likelihood model also recovered the main components of the generative process. Estimated smooths matched the true response functions to covariates (Figure 3; cf. Appendix S1: Figure S2).

**Figure 3.**
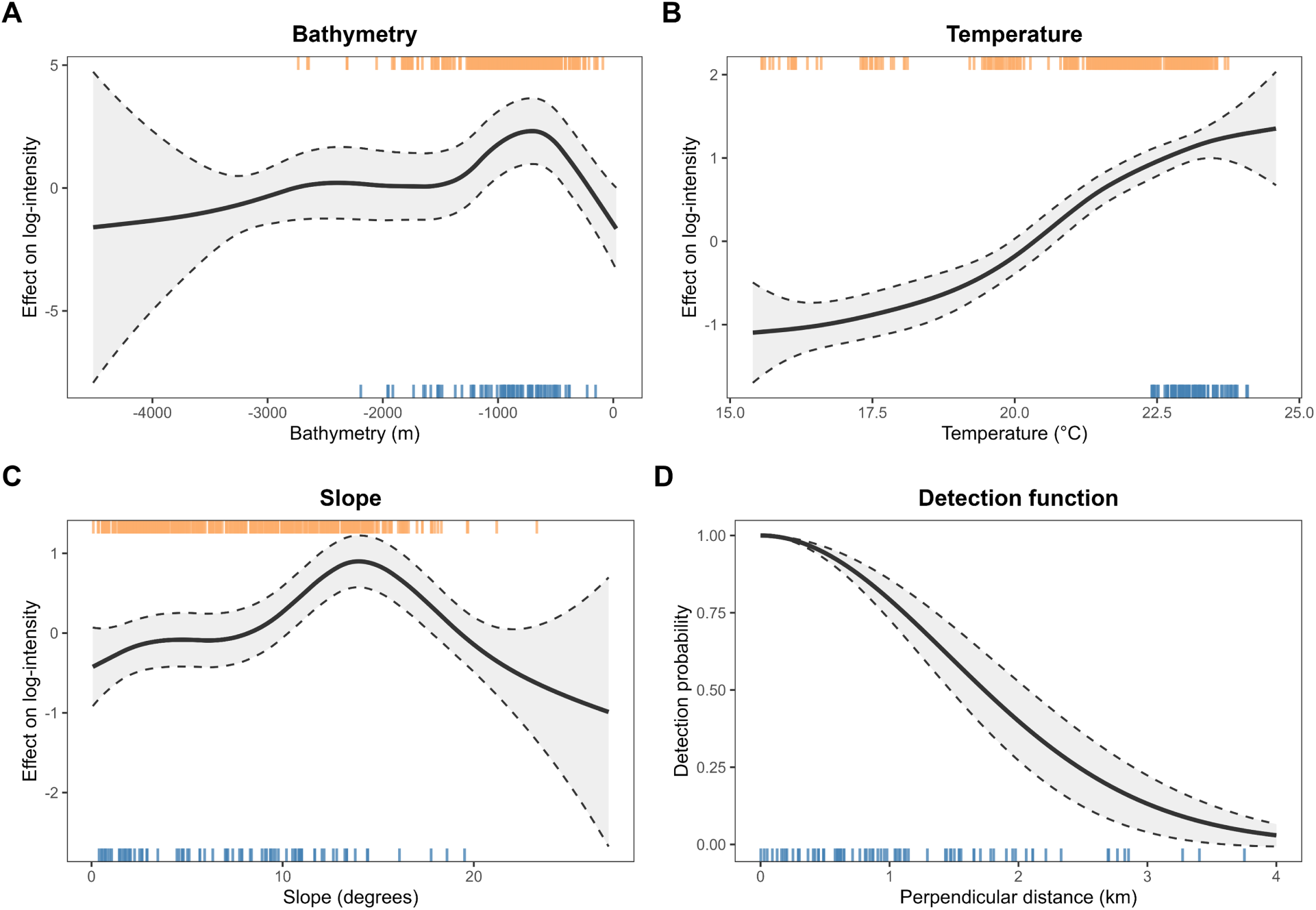
Partial effects of **(A)** bathymetry, **(B)** temperature, **(C)** slope on the log-intensity, and **(D)** the survey detection function. Solid lines are posterior means; shaded bands are 95% credible intervals from 2,000 posterior samples. Covariates were modelled on scaled values but are plotted in raw units for interpretability. Effects were evaluated over the covariate space within the inner boundary (model projection domain). Rugs show data support (animals sighted) by dataset: top orange ticks = whale watching (WW), blue ticks = survey. Panel (D) shows the fitted half-normal detection function over perpendicular distance; bottom ticks indicate observed distances.

Bathymetry showed a low–mid-depth optimum matching the Gaussian peak used in the generative model at 750 m. Temperature increased across the observed range, consistent with the simulated logistic rise in suitability; note that WW contributed most of the dynamic support at lower values (orange rugs). Slope displayed a peak near the simulated optimum (15^◦^), with a smaller effect size than bathymetry and wider intervals at steep values where data are scarce. Model extrapolation in covariate space was limited and occurred mainly at the deep end of the bathymetry range (see Appendix S1: Figure S12). The fitted August detection curve showed the expected half-normal decline with perpendicular distance. The posterior mean of the scale parameter was *σ* = 1.48 km (95% CrI: 1.23 km–1.77 km). For reference, the simulation used a half-normal with *σ* = 1.63 km, which lies well within the credible interval.

#### (iii) Absolute abundances

Overall, the integrated model recovered monthly absolute group abundance well across the year. The mean signed relative error was +10.7%. Six months were within 10% of the simulated mean, including the seasonal peak (July to September), where errors were small (−6.8% to −2.7%). The largest positive errors were in December (25.0%) and June (23.0%). In every month, the simulated means lay well within the model’s corresponding 95% credible interval.

### 3.2 Multi-scenario comparison

Overall, the integrated model achieved the highest spatial recovery (median Spearman *ρ* = 0.824, IQR 0.765–0.877), followed by the survey-only model (*ρ* = 0.750, IQR 0.659–0.823) and the WW-only model (*ρ* = 0.668, IQR 0.532–0.798). The integrated model outperformed the WW-only model in 91.6% of month-specific comparisons (median Δ*ρ* = 0.134) and the survey-only model in 89.0% of comparisons (median Δ*ρ* = 0.064). The two gains showed contrasting sensitivity to the covariate–field balance. Gains over the WW-only model increased as the point-generating surface became more strongly field-driven (C1S3 > C2S2 > C3S1) and were largest at the shortest GRF range (*ρ*_water_ = 100 km; median Δ*ρ* = 0.260- 0.266 for C2S2 and C1S3), consistent with the WW-only model’s difficulty recovering fine-scale spatial structure. Gains over the survey-only model showed the opposite pattern, being largest in covariate-dominated scenarios (C3S1 > C2S2 > C1S3) and across all ranges (median Δ*ρ* = 0.093–0.145 for C3S1), reflecting the survey model’s reliance on a single August transect and its reduced capacity to extrapolate covariate effects temporally. In field-dominated scenarios, where the spatial pattern is largely static across months, the survey model’s August snapshot transferred more reliably, narrowing the gap with the integrated model (Figure 4).

**Figure 4.**
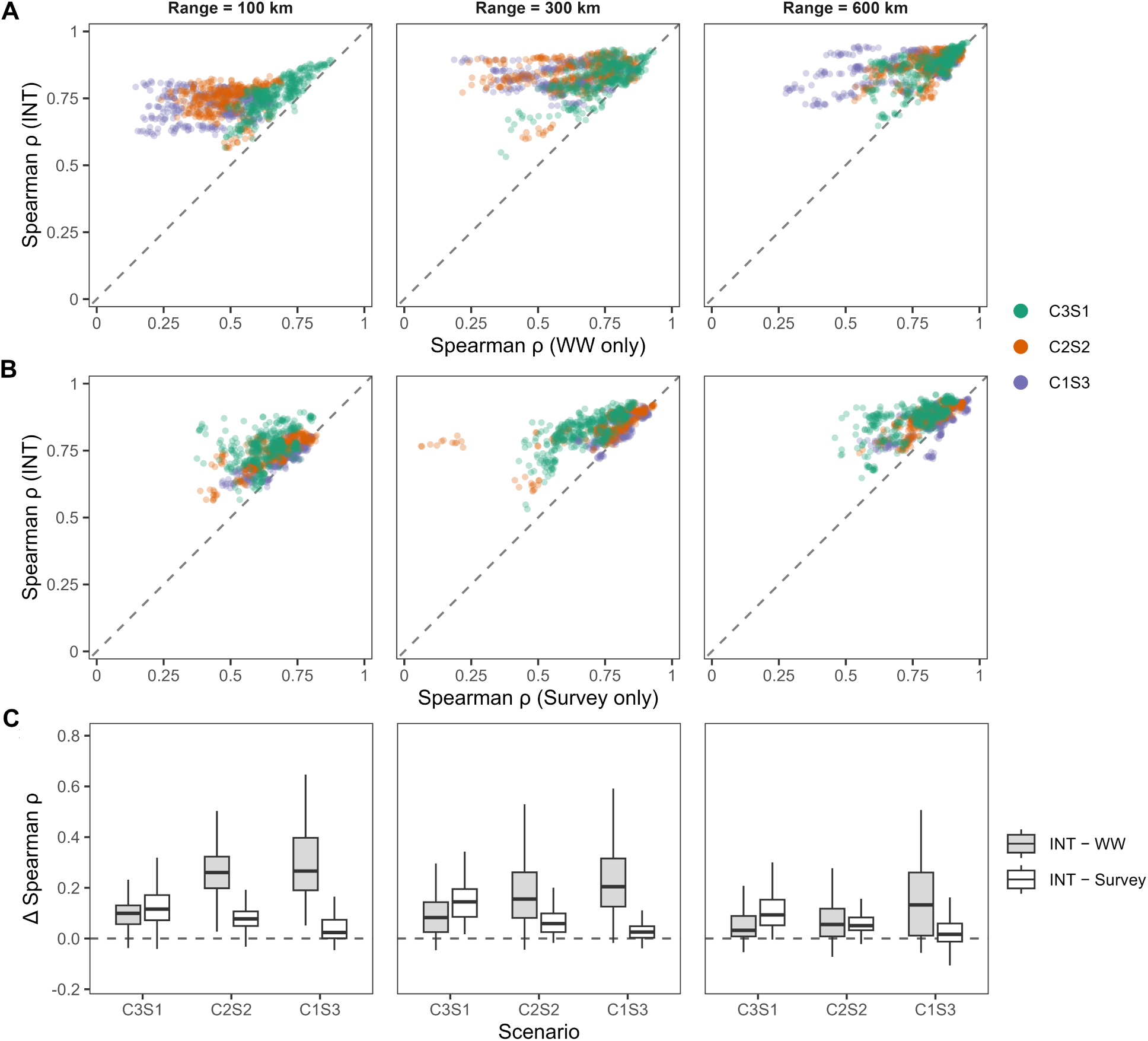
Spatial recovery performance across model types and simulation scenarios. **(A, B)** Month-specific Spearman rank correlations between the simulated habitat-suitability surface and the predicted log-density surface of the integrated model versus the WW-only **(A)** and survey-only **(B)** models. Each point represents one month–replicate combination; colours indicate the covariate–field balance (C3S1: covariate-dominated; C2S2: equal; C1S3: field-dominated); the dashed diagonal denotes equal performance. **(C)** Gain from integration (Δ*ρ*) over the WW-only (grey) and survey-only (white) models, summarised as boxplots per scenario; the dashed line indicates Δ*ρ* = 0. Facets correspond to the simulated GRF range (*ρ*_water_ ∈ {100, 300, 600} km).

### 3.3 Case study

Figure 5 A shows that the GRF explained most of the variation in the linear predictor: the combined surface closely followed the GRF, indicating that broad-scale spatial structure dominated the fitted pattern. Within the covariate block, slope had a positive association with log-intensity. Temperature showed a shallow hump-shaped response with a maximum at intermediate values. Chlorophyll displayed a negative association with log-intensity across most of its range. Effort coverage across covariate space (Appendix S1: Figure S13) showed that extrapolation was concentrated in the upper range of chlorophyll and slope (high concentrations and steep slopes), which were weakly sampled by both data sources. Temperature was well constrained across mid-range values, with only limited extrapolation at the coolest and warmest extremes. Because temporal variation entered the model only through the dynamic covariates, and their amplitudes were modest, temporal fluctuations in the spatial predictions were small; see Appendix S1: Figure S14.

**Figure 5.**
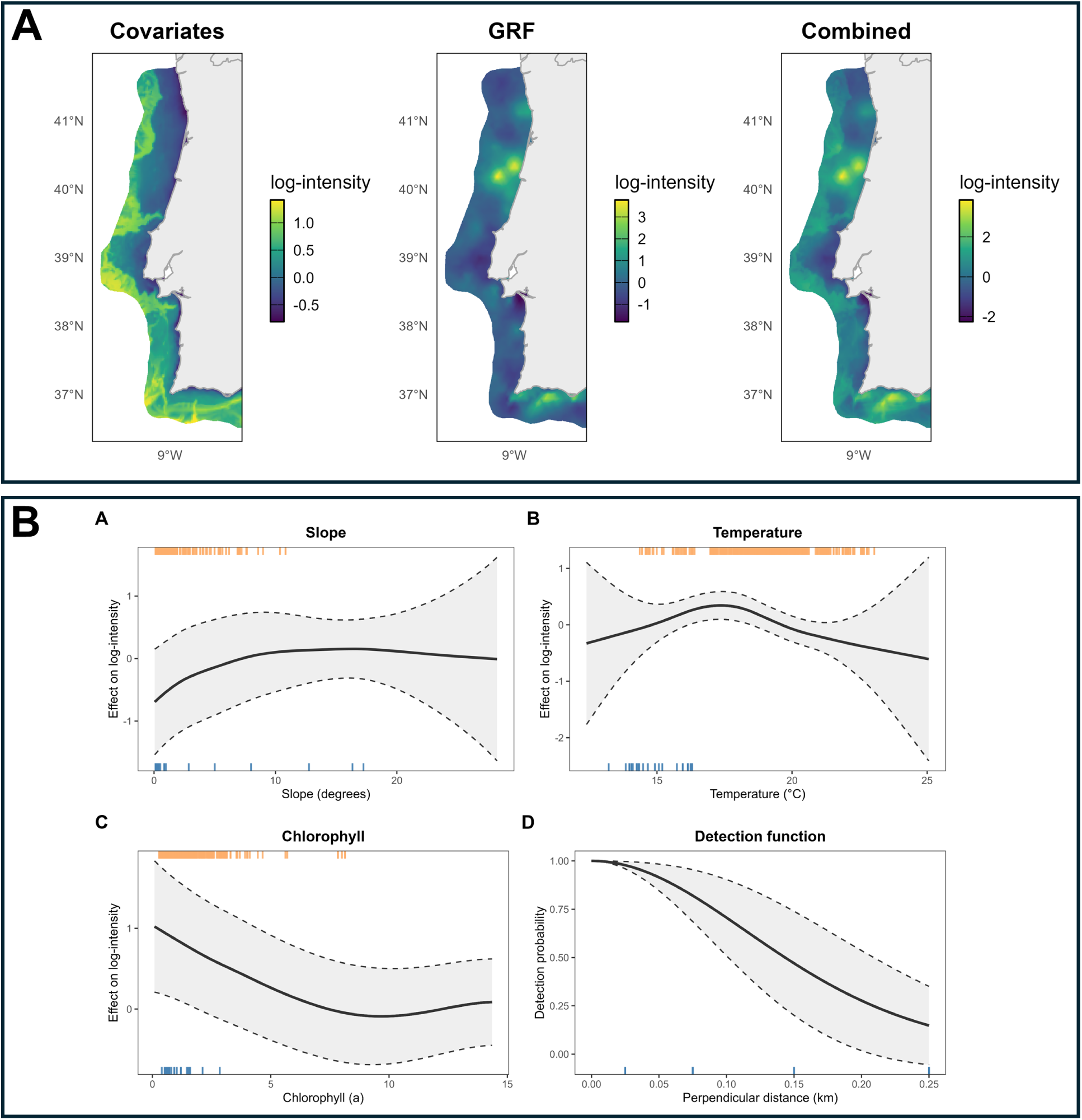
Posterior outputs for the common dolphin case study. **(A)** February (survey month) component maps from the integrated model: covariate-only contribution; the Gaussian random field (GRF), *ω* **s**; and the combined linear predictor. **(B)** Partial effects: posterior mean (solid) with 95% credible bands (shaded) for the covariates, and the fitted half-normal detection curve for the survey data. Covariate effects were estimated on standardised values but are plotted in raw units for interpretability. Rugs indicate data support by dataset: top orange ticks = pooled whale watching (WW); bottom blue ticks = SPEA survey (for detection, rugs show observed perpendicular distances).

## 4 Discussion

Our study was motivated by a common constraint in dynamic species distribution modelling: available data often arrive as puzzle pieces, observed through different lenses and sampled unevenly across space and time. A joint point-process view aligns those pieces to a common ecological state by linking observation-specific likelihoods to a shared latent intensity surface. We used cetaceans as a motivating case because the space–time data mismatch is particularly acute in dynamic marine systems and has clear management relevance, but the contribution is general. In our formulation, designed surveys anchor absolute scale and broad spatial structure, while opportunistic records contribute temporal replication within limited, biased footprints.

In idealised simulations, this complementary role of the two data sources was reflected in model performance: integration captured covariate-driven temporal variation from opportunistic data while preserving the large-scale spatial structure anchored by the survey. Across the scenario grid, integration improved recovery in most cases, with gains over the WW-only model largest when residual spatial structure was strong and fine-scale, and gains over the survey-only model largest in covariate-dominated scenarios, where a single August transect alone had limited capacity to resolve temporal change. At the same time, the case study highlights the constraint of sparse survey replication. When covariate effects are weak and residual spatial structure dominates, temporal variation in predictions remains modest because dynamic change can only enter through covariates learned mainly within opportunistic footprints. These results therefore clarify both the potential and the limits of spatiotemporal integration under uneven coverage, and make explicit what additional survey replication would buy in terms of identifiable spatiotemporal residual structure. Below, we outline the underlying assumptions and trade-offs.

### 4.1 Spatiotemporal structure with uneven effort

In our model we specified a time-invariant spatial GRF, *ω*(**s**), and allowed temporal change only through dynamic covariates *z _j_* (**s**, *t*) in the shared linear predictor. This specification followed from the data configuration: a single month of broad-coverage survey with few detections, paired with small coastal opportunistic footprints across months. Under such support, a space-time GRF, *ω*(**s**, *t*), is essentially unidentifiable outside the WW footprints; it would learn month-specific residual structure almost exclusively inside those areas and extrapolate it elsewhere with little support. Temporal variation was therefore learnt from the time-varying covariates, which were well supported within opportunistic footprints. Outside those footprints, month-to-month differences were driven by the covariates together with the broad spatial pattern encoded by *ω*(**s**). This design carries two explicit assumptions: first, that the broad spatial pattern is constant throughout the year; and second, that covariate responses learnt within opportunistic footprints transfer to unsampled areas, so temporal change outside footprints is governed by the covariates.

While this setup performed well in simulation, the case study showed a weaker covariate signal. Three features explain this. First, survey detections were few, which limited power to resolve covariate–response relationships; credible intervals were wide, consistent with a small sample size and limited effort. Second, the survey was conducted in a single month and therefore spanned only a narrow range of the dynamic covariates (SST and chlorophyll-*a*); estimation of these responses relied largely on WW records within coastal footprints. Third, there was residual spatial structure not captured by the covariates, which the stationary GRF absorbed as intended. Together, these features meant that the GRF contributed most of the spatial structure and therefore month-to-month differences in predictions were small. We therefore interpret covariate effects in the case study primarily as descriptive associations under limited support, rather than as robust causal signals. We explored two alternatives for modelling temporal structure and did not retain them. Adding a smooth month effect confounded with the dynamic covariates and absorbed their signal. Decomposing the latent field into a persistent spatial term plus a month-specific residual term yielded a residual that was estimable only within WW footprints and again confounded with the covariates. We therefore retained the simpler spatial GRF specification. Although the assumptions of our model are strong, the single-survey configuration reflects common data availability and makes the trade-offs explicit. We therefore retain this specification to illustrate its limitations as well as its utility. Additional survey months in this system would justify a full spatiotemporal GRF with identifiable temporal correlation and would directly improve identifiability of both covariate responses and residual spatiotemporal structure.

### 4.2 Preferential sampling

Our WW data are a classic case of effort-biased and preferential sampling. By definition, these biases occur when observation effort is stochastically related to the underlying ecological state, such that sampling locations or durations are more likely where true density is high (Diggle et al. 2010). Conditioning the likelihood on sampler footprints (month-specific availability polygons *A*(*t*) for WW) restricts integration to areas actually visited, correcting outside-footprint availability bias and preventing spurious extrapolation. However, even with temporally varying *A*(*t*), true effort might be uneven within the footprint; residual within-footprint preference may persist. Converting available GPS tracks into an effort surface would require behaviour-state segmentation and assumptions about detectability that could dominate the offset in our small coastal study areas, so we did not pursue this. Other temporally rich opportunistic datasets (e.g., fisheries data, structured citizen-science schemes, or camera-trap networks) can be incorporated within the same framework, but may be less affected by this limitation when effort and detectability are available. In such cases, observation models can include explicit effort or detection terms, reducing residual preferential-sampling bias and making the opportunistic contribution more robust.

### 4.3 Conclusions and outlook

Several extensions to this work are both feasible and worthwhile exploring. First, additional survey months would justify a fully spatiotemporal random field with identifiable temporal correlation; quantifying how many surveys are needed for robust space-time GRF estimation is a next step. Second, moving toward animal density might be achieved with marked point-process formulations for group size. Third, extending this framework to other survey modes opens promising research avenues. A variety of survey modes contain information about species distributions and animal density, but resolve space and time in fundamentally different ways. These include passive acoustic monitoring, drone-based surveys, tag-based animal-movement data, environmental DNA, and satellite detections (e.g. Marques et al. 2013; Fretwell et al. 2014). Incorporating such data through dedicated observation-process likelihoods, where appropriate, would further broaden the applicability of the framework.

In all these settings, integrated models do not substitute for well-designed surveys, whose value remains fundamental, but they can increase the information gained from each observation mode. Used together, structured surveys and additional data streams—in particular, though not exclusively, opportunistic sources as illustrated here—can yield more reliable, consistent, and temporally resolved predictions than dedicated surveys alone, improving the utility of density surfaces for conservation planning and management.

## Acknowledgments

The project that gave rise to these results received the support of a fellowship from the ‘la Caixa’ Foundation (ID 100010434) to MK. The fellowship code is (LCF/BQ/DI23/11990054). The Atlantic Whale Deal project (EAPA_0004/2022), cofunded by the Interreg Atlantic Area 2021–2027 programme, supported the acquisition of computing equipment for data processing. FA and MF had the support of Fundação para a Ciência e a Tecnologia (FCT) throughout the strategic projects UIDB/04292/2025 (https://doi.org/10.54499/UID/04292/2025) granted to MARE and LA/P/0069/2020 (https://doi.org/10.54499/LA/P/0069/2020) granted to the Associate Laboratory ARNET. FA was also supported by an FCT research contract under the Scientific Employment Stimulus Call (https://doi.org/10.54499/2023.08949.CEECIND/CP2851/CT0001). MF was financially supported by the Atlantic Whale Deal project. TAM acknowledges the US Navy funded AMERICANO project. MPM, MK and TAM thank partial support by CEAUL (funded by FCT, Portugal: https://doi.org/10.54499/UIDB/00006/2020). We thank all whale-watching partners and survey teams for data collection and support.

## Author Contributions

MK, MF, and TAM conceived the study and designed the methodology, with additional conceptual input from FL, LT, and VMP; MK led the writing of the original draft and performed all analyses with input from FL, LT, and VMP; data were curated by MK, FA, MPM, NO, JC, FM, SM, and AR; supervision was provided by FA, FL, MF, and TAM; all authors contributed to review and editing and approved the final version for publication.

## Appendix S1: Supporting Information

### Section S1: Uncertainty propagation and model assessment

Uncertainty enters the model through the survey detection component *g*(*d*; *σ*) and through the shared latent density surface *λ*(**s**, *t*). In traditional two-stage density surface modelling, detectability is typically estimated first and then treated as a fixed offset in the spatial model, so uncertainty in *g*(*d*; *σ*) is not carried forward and downstream uncertainty can be underestimated. Approaches to propagate this uncertainty have been developed recently (Bravington et al. 2021), but challenges remain if individual level covariates are present. By contrast, in the specified one-stage formulation, detectability and the latent density are estimated jointly, retaining posterior dependence between model components. As a result, uncertainty propagates directly to any quantity computed from the joint posterior, such as predicted density surfaces, abundance summaries, or covariate response functions. Posterior uncertainty for such quantities can be obtained directly from posterior samples of the joint model. For model diagnostics, we focused on checks that respect the joint-likelihood structure and the differing sampling domains of the two data sources. For the line-transect component, we assessed the detection submodel using posterior-predictive binned counts of perpendicular distances and checked sensitivity to the detection-key specification (Figure S8). For abundance, we additionally compared a plug-in approximation to the full posterior predictive distribution (Figure S9). For model checking, we relied on controlled simulations as the primary diagnostic, because the point-generating surfaces are known.

### Section S2: Covariate smooths

We use smooth *f _j_* (·) terms rather than linear effects because species responses to environmental covariates are rarely linear. Each *f _j_* is modelled as a 1-D Matérn Gaussian field over the covariate domain, using an SPDE representation again with PC priors. Following the SPDE formulation, we discretise *f _j_* (·) on a 1-D mesh over the observed covariate range and use a finite-element basis of degree 2 (piecewise-quadratic functions):

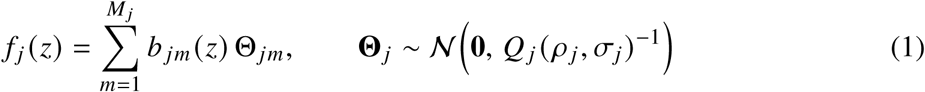

where *z* is the covariate value, *M_j_* is the number of (mesh) nodes in the 1-D mesh for covariate *j*, *m* indexes the mesh nodes; *b _jm_* (*z*) are the quadratic finite-element basis (shape) functions, Θ*_jm_* are the Gaussian weights at node *m*, Θ*_j_* = (Θ*_j_*_1_, … , Θ*_j Mj_*)^⊤^ is the weight vector; and *Q _j_* ( *ρ _j_*, *σ_j_*) is the SPDE precision matrix parameterised by the practical range *ρ _j_* and marginal standard deviation *σ_j_* . While these non-linear smooths are well-suited for capturing complex habitat preferences, they are a flexible option rather than a requirement for the core data integration framework. Each *f _j_* uses the same priors across all analyses: a PC prior on the marginal SD with ℙ(*σ* > 1) = 0.05; the range *ρ _j_* is fixed to the covariate span to encourage smoothness and mitigate overfitting.

### Section S3: Detailed simulation methods

#### Occurrence generation

We generated animal groups with the virtualspecies R package (Leroy et al. 2016), using the region around the Azores (central North Atlantic, Portugal) as our study system. The virtual species was loosely inspired by short-finned pilot whales (*Globicephala macrorhynchus*), a relatively common delphinid in the region during the summer months (Alves et al. 2019). The group distribution was defined by three environmental covariates. These included two static layers (bathymetry and seafloor slope) and one dynamic layer (sea-surface temperature, SST). All layers considered the same 2 km marine grid and were masked over land; SST was averaged to monthly means for 2019. We specified a logistic increase of suitability with SST and Gaussian peaks centred at 750 m depth and 15° slope. Additionally, we considered a GRF to represent the unmeasured spatial structure in occurrences that always occurs in real data. For that, we simulated a broad-scale residual GRF (Matérn via a barrier SPDE) to mimic a persistent, unmeasured large-scale pattern (e.g. a large-scale productivity or front structure not captured by our covariates; Figure S1).

**Figure S1.**
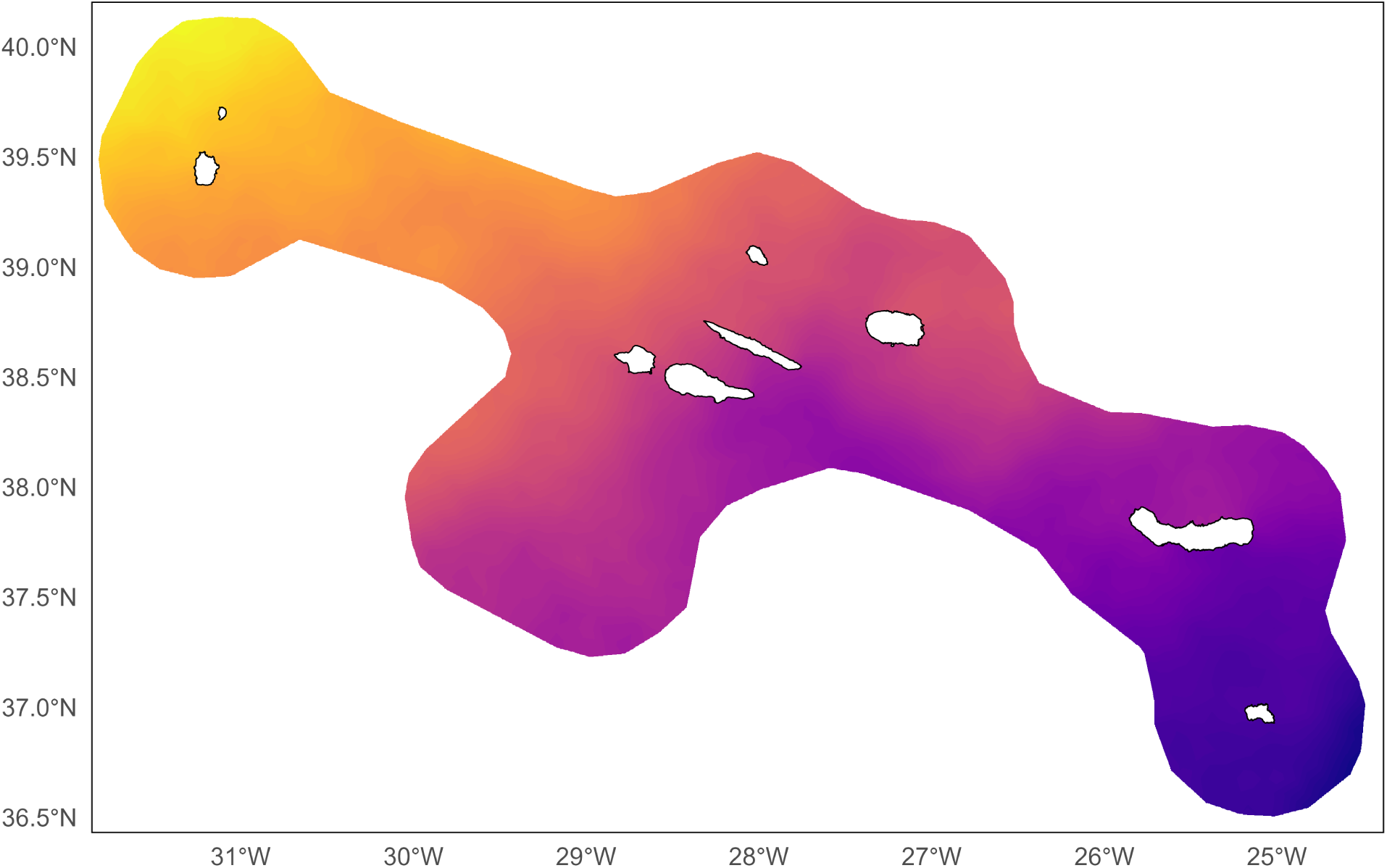
The realisation of the barrier-aware Matérn SPDE used to generate the residual spatial structure in the simulation.

We set a long practical range over water (*ρ* = 600 km) and suppressed correlation across land by shrinking the range on barrier elements (see Bakka et al. (2019)). We simulated this field on the same triangulated mesh (a finite-element discretisation of space) used for modelling, projected it onto the 2 km grid, and combined it additively with the covariate terms to produce the final suitability surface. Specifically, higher suitability scaled linearly with higher density field values, alongside the SST (logistic) and the depth and slope (Gaussian) terms (Figure S2).

**Figure S2.**
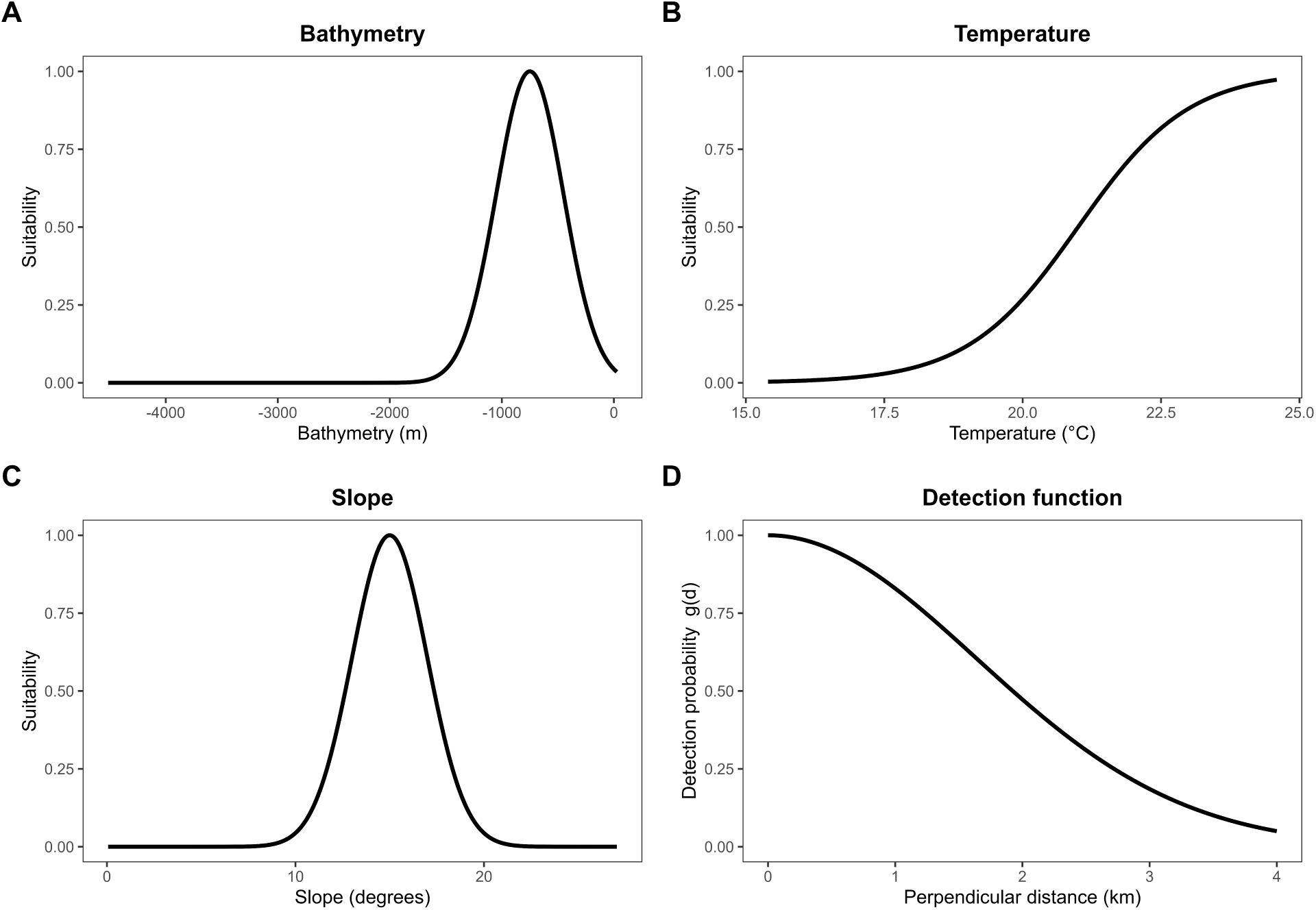
Suitability curves used in simulation, plotted against covariate values: (A) Bathymetry (m), (B) Temperature (°C), (C) Slope (degrees), and (D) half-normal detection function vs perpendicular distance (km).

All covariates contributed additively with equal weight, and the resulting suitability was then scaled to [0, 1]. We converted monthly suitability to occurrence probability using a logistic link with *γ*_1_ = −0.05 as the steepness parameter and *γ*_2_ = 0.5 as the inflection point. Presences were generated by treating each grid cell as a Bernoulli trial with success probability equal to its occurrence probability. To set realistic abundances, we targeted 0.4 groups/100 km^2^ (Fernandez et al. 2022). Multiplying this rate by the total marine area of the 2 km grid gave a reasonable number of groups. We then calibrated the overall sample size so that the number of simulated presences in the peak month (August) matched this expectation. For each month, we generated 30 independent daily realisations across the region to emulate day-to-day movement. Because occurrences were grid-based, initial locations fell at cell centroids; for point-process inputs, we jittered each point within its cell to assign unique coordinates without changing the underlying probabilities. Monthly point-generating surfaces and one representative day of simulated occurrences are shown in Figure S3.

**Figure S3.**
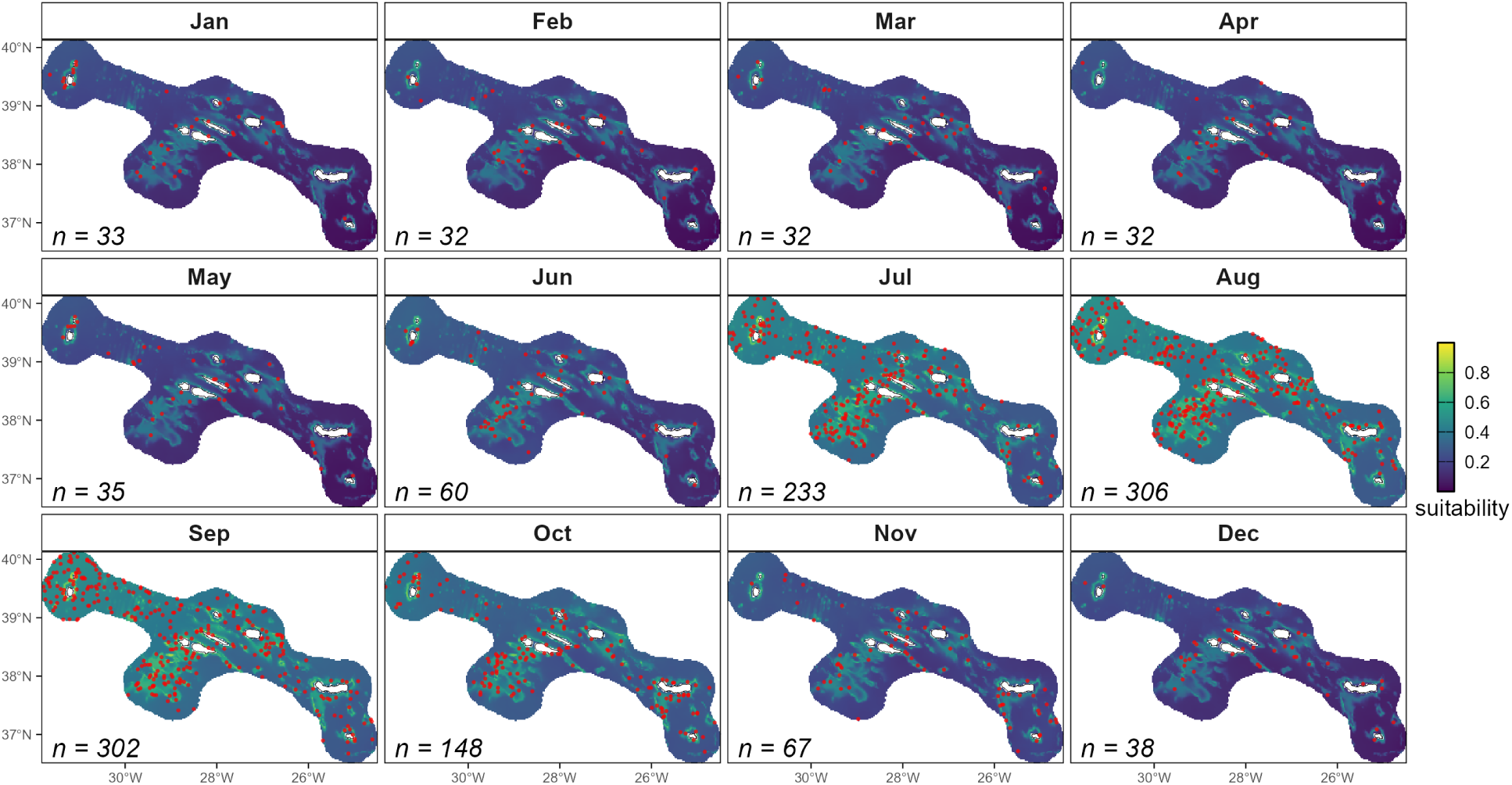
Monthly habitat suitability (0–1) for the Azores, used as the point-generating surface in the simulation. One randomly selected day of occurrences per month is overlaid (red points). The label “n = … ” reports the mean daily number of occurrences across the 30 simulated days in that month.

##### (i) Line-transect survey

We used the equally spaced zigzag survey layout and strata from Faustino et al. (2010) for the Azores, taking their track lines as the sampling frame. Because shipboard cetacean surveys in the region are typically scheduled for summer when sea state is calmer and daylight is longer, we simulated survey-derived detections for August. To tie the survey to the latent region-wide occurrences, we split the track lines into daily segments assuming 12 h on-effort at 11 kn, yielding up to 200 km per day and completing the design in 29 days. For each survey day we buffered the lines by 4 km on either side (the truncation distance) to define availability, intersected the buffer with the respective day’s simulated presences, and kept only groups inside the truncation distance. For every retained group we computed the perpendicular distance to the nearest same-day track line and assigned a half-normal detection probability parameterised so that *g*(4 km) = 0.05 (*σ* = 1.63 km). For simplicity we assumed perfect detection on the line (*g*(0) = 1) and let detectability depend only on perpendicular distance, with no effects from sea state or other covariates. We then applied Bernoulli thinning with that detection probability. Across ∼4963 km of on-effort track line in August, half-normal thinning reduced 165 available groups to 79 detections, which we used for modelling (Figure S4).

**Figure S4.**
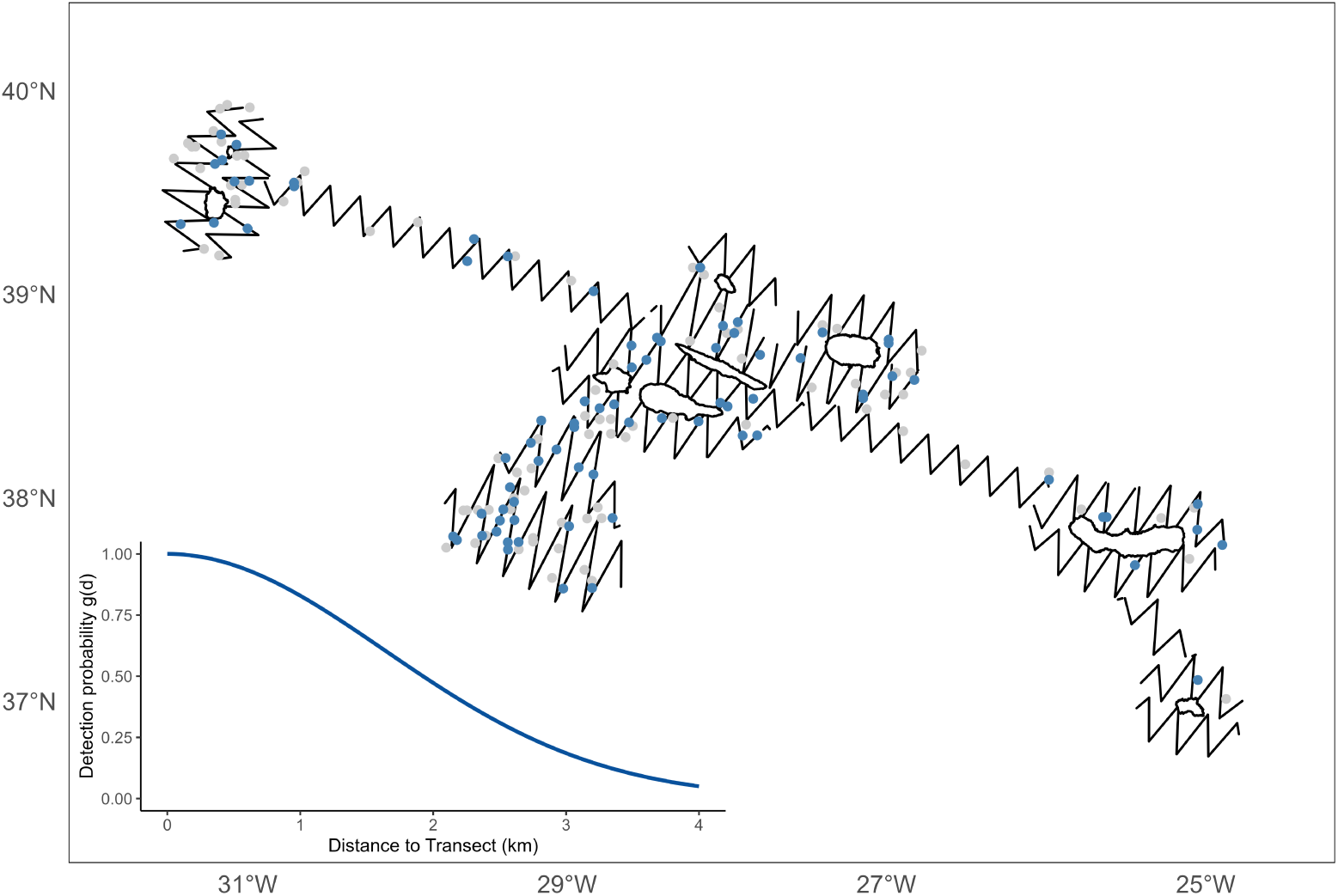
Transect lines from the virtual survey across the Azores EEZ (black). Points show all simulated groups within the truncation distance (4 km; *n* = 165); blue points are retained after thinning by a half-normal detection function (*g* 4 km = 0.05, *σ* = 1.63 km; *n* = 79). Inset: the half-normal detection curve used for thinning, truncated at 4 km.

##### (ii) Whale watching

To avoid arbitrary effort polygons, we derived monthly availability directly from WW sightings in the region. Specifically, we obtained WW effort with MONICET (González García et al. 2023), an open citizen-science platform that has collected georeferenced sightings in the Azores since 2009 (records downloaded via the Global Biodiversity Information Facility (GBIF 2025)). For each month, convex hulls were built from the corresponding MONICET locations and clipped to sea. Within the polygons, we assumed constant detection; operationally, in the sampling of our simulated data we intersected the group locations with the month’s polygons and treated all group locations inside as available to WW, while group locations outside were unavailable. Since operators are usually busiest in summer and sporadic in winter, we simulated this temporal effort as increasing by one day per month to a July peak (7 days), then declining symmetrically to December (2 days). For each month, the active days were drawn at random (without replacement) from the calendar days. Because of the spatiotemporal effort and the fact that the virtual species preferred warmer temperatures, the number of simulated occurrences captured by WW varied seasonally: August showed the highest count within the WW effort area (*n* = 163), whereas January recorded only one detection (*n* = 1; Figure S5, Table S3).

**Figure S5.**
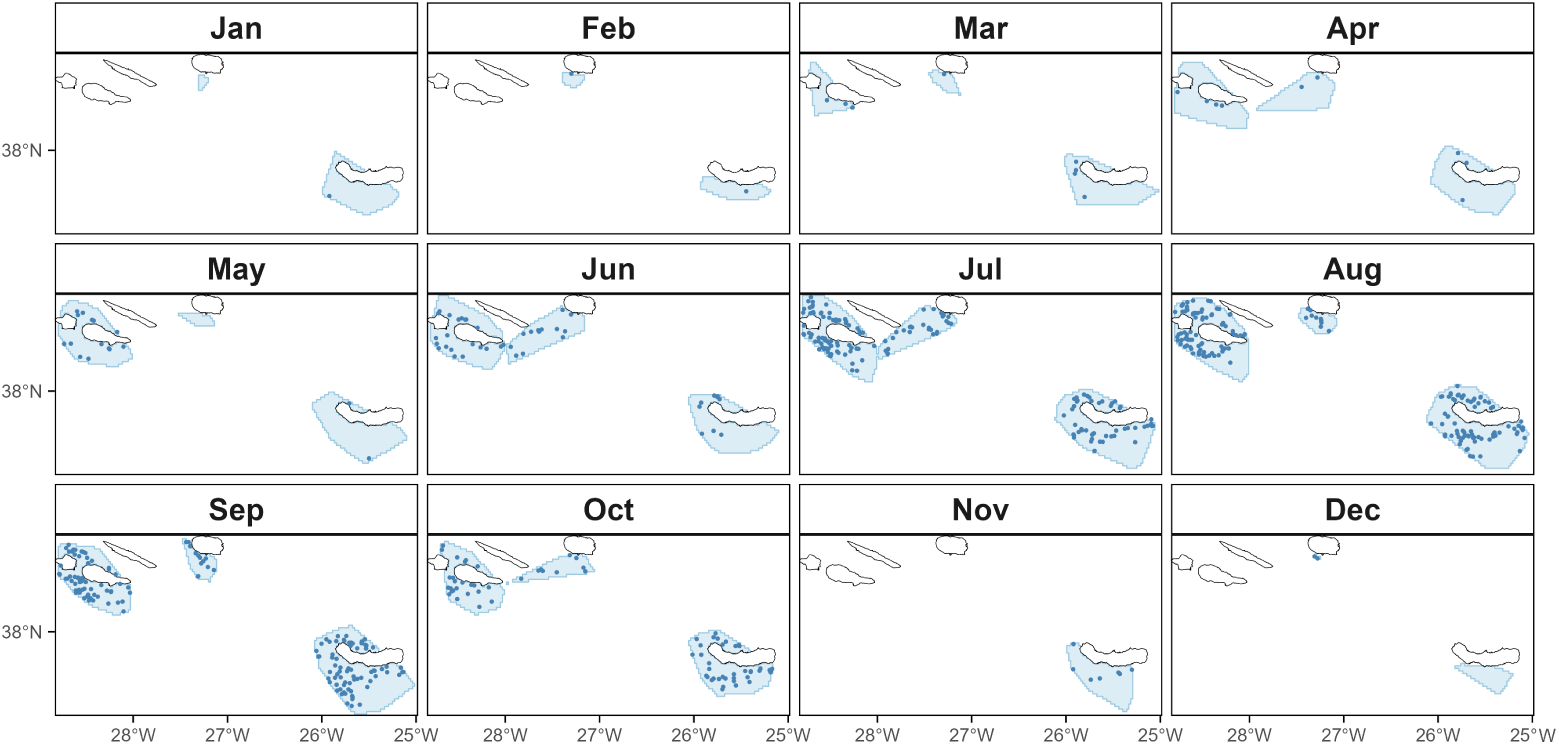
Monthly availability polygons (light blue) derived from MONICET whale-watching (WW) records. Dark blue points are the virtual WW sightings used in the simulation. The western group (Flores and Corvo) is omitted because there is no WW effort.

#### Modelling

We built the spatial field on a 2-D triangulated mesh in an equal-area Azores projection. The model domain was a boundary polygon ∼25 km out from the survey track lines and an outer polygon ∼150 km beyond that to buffer boundary effects. To control resolution, we seeded vertices with hexagon lattices: 2 km over islands and 8 km over the inner hull (Figure S6). To prevent correlation across land, we tagged mesh triangles with centroids on land, which were later used to define the barrier SPDE (Bakka et al. 2019).

**Figure S6.**
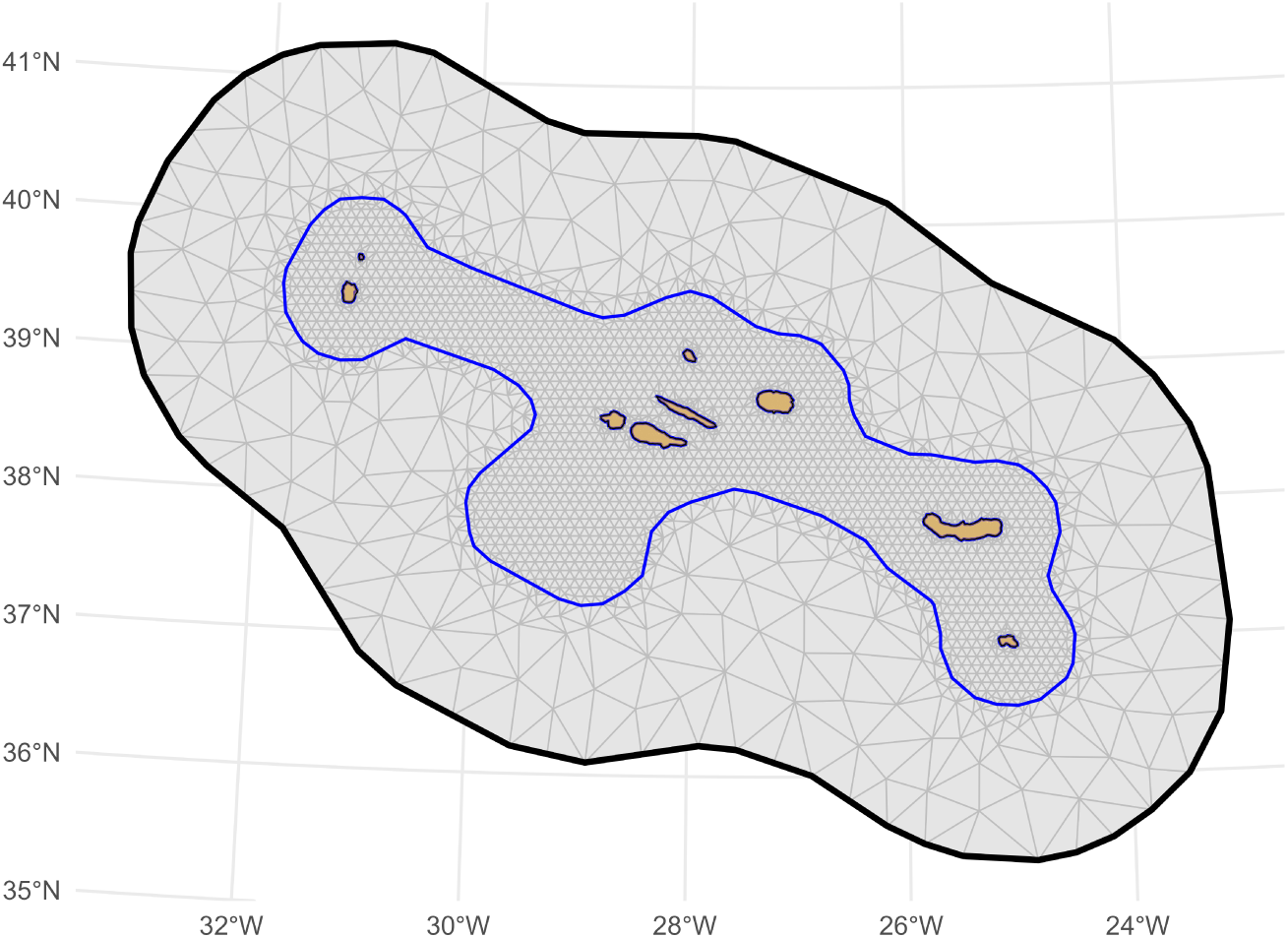
SPDE mesh for the Azores domain. The triangulated mesh defines the 2-D spatial field. Orange elements are mesh triangles whose centroids fall on land and are treated as barrier triangles to suppress correlation across land (barrier SPDE).

The at-sea correlation range was fixed to match the simulation scale, and we put a PC prior on the marginal standard deviation with a 5 % tail above 1. To suppress correlation across land, we reduced the range on barrier elements to 1 % of the at-sea range, which effectively prevented dependence from propagating through islands while preserving continuity at sea. The barrier model was then used as the spatial field in the LGCP. Besides the spatial field, the latent density included the same covariates that we used for simulating the data: bathymetry, slope, and monthly SST. All covariates were standardised and entered into the model as 1-D SPDE smoothers using a quadratic (degree-2) finite-element basis on a regular mesh spanning the observed covariate range (see Section S2). We fitted a single joint model with a shared latent predictor and two likelihoods. Numerical integration of the point process likelihoods was performed on a finer mesh obtained by subdividing the SPDE mesh, giving 2 km quadrature spacing to match the covariate resolution, while the GRF used the coarser base mesh for computational efficiency. Because the simulation supplied the ground truth, we evaluated recovery by comparison of the posterior mean log group density maps with the simulated point generating suitability maps. Besides visual comparison, for each month we calculated Spearman’s rank correlation across all valid grid cells between simulated suitability and the model’s posterior means. We also evaluated our model by checking component posteriors such as covariate smooths, the spatial field, and the estimated distance detection function. Finally, we estimated monthly absolute group abundance from the integrated model, and compared these estimates with the simulated abundances.

### Section S4: Additional tables and figures

**Table S1:** Abbreviations used in the main text.

| Abbreviation | Meaning |
| --- | --- |
| CI | Confidence interval |
| CrI | Credible interval |
| GMRF | Gaussian Markov random field |
| GRF | Gaussian random field |
| INLA | Integrated nested Laplace approximation |
| IPP | Inhomogeneous Poisson process |
| LGCP | Log-Gaussian Cox process |
| LOOCV | Leave-one-out cross-validation |
| MCMC | Markov chain Monte Carlo |
| PC prior | Penalised complexity prior |
| SDM | Species distribution model |
| SPDE | Stochastic partial differential equation |
| WAIC | Watanabe–Akaike information criterion |

**Table S2:**
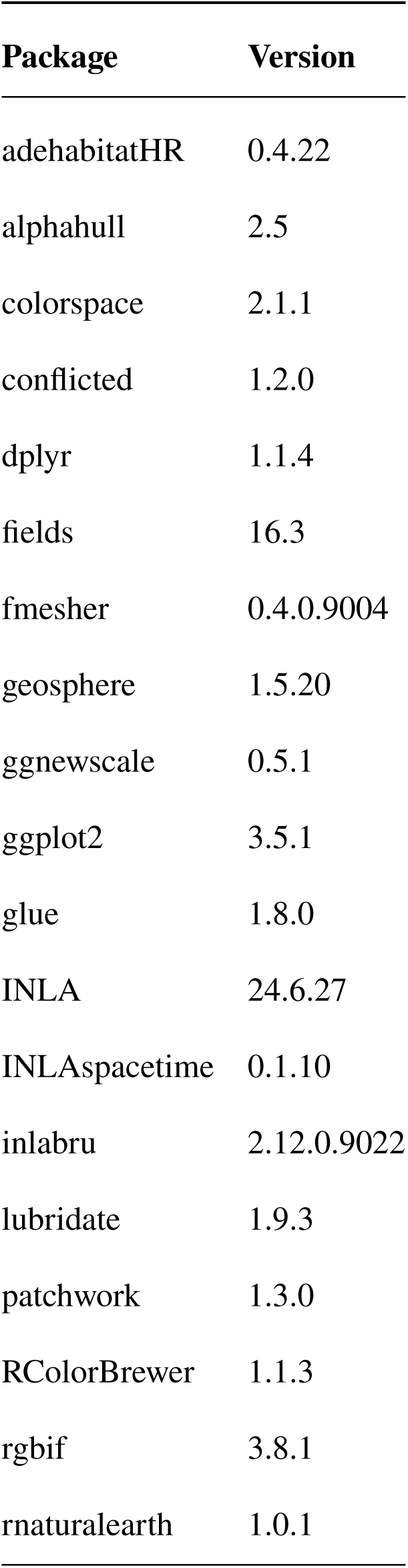
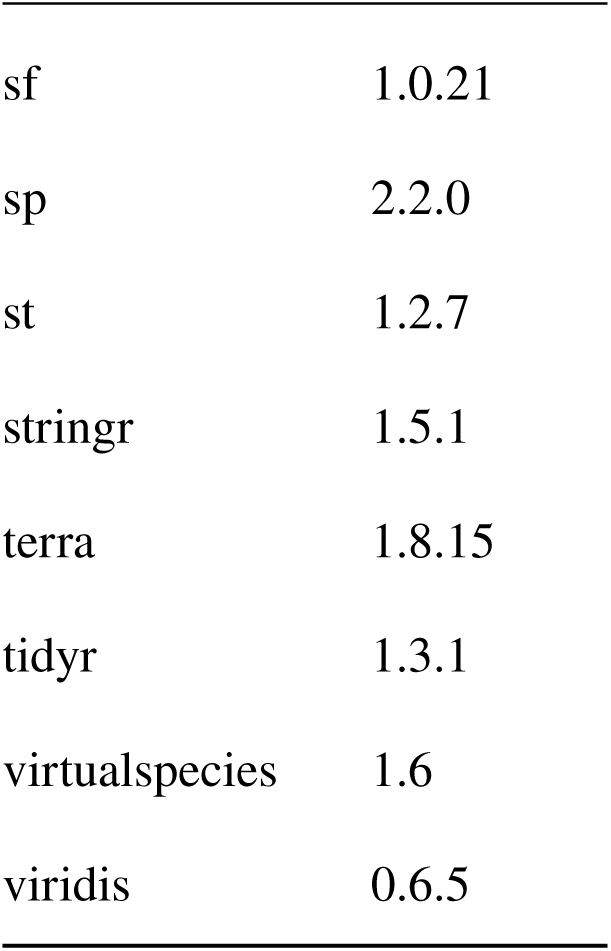
R packages used in the analyses (versions at analysis time).

**Table S3:**
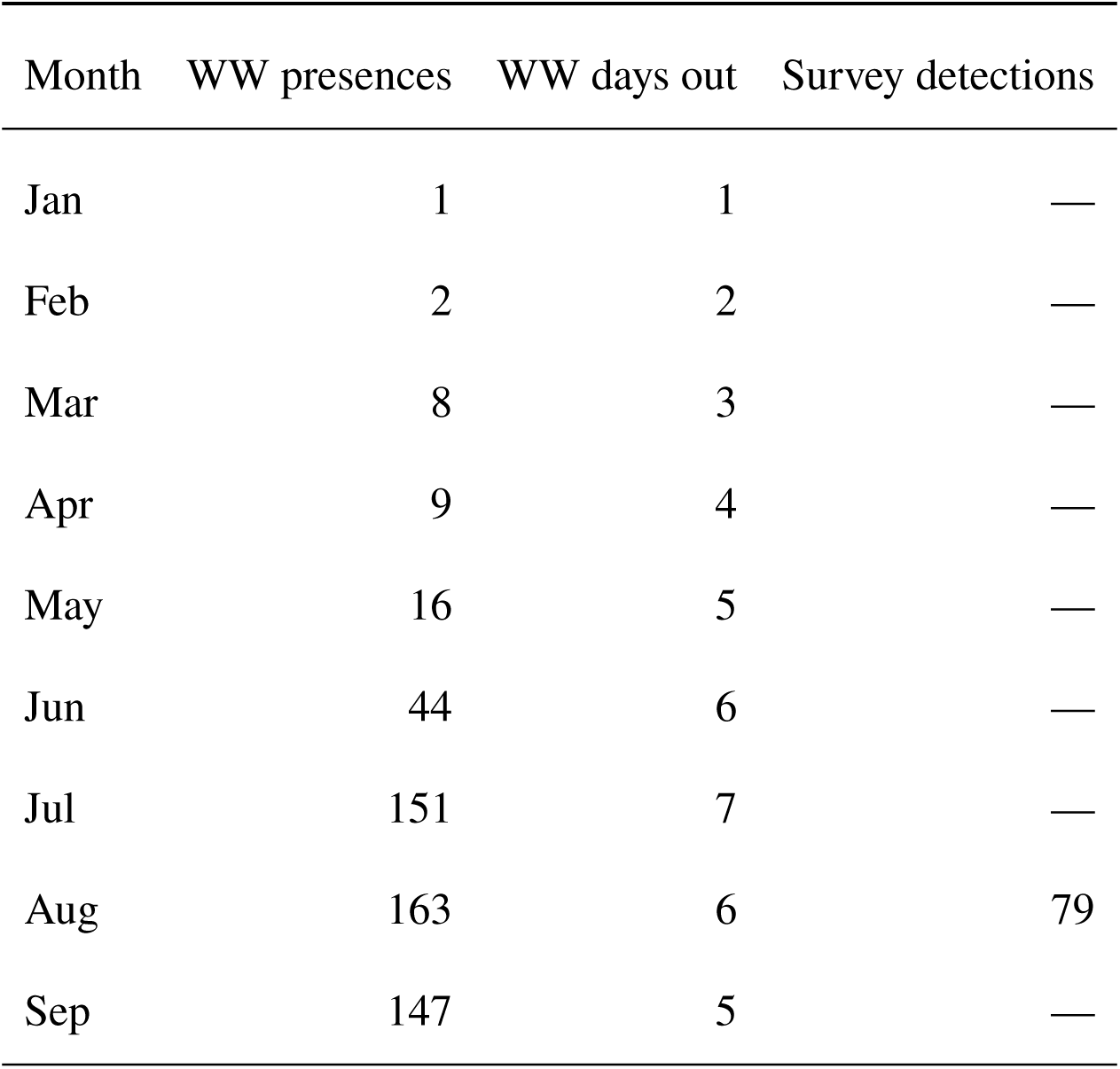
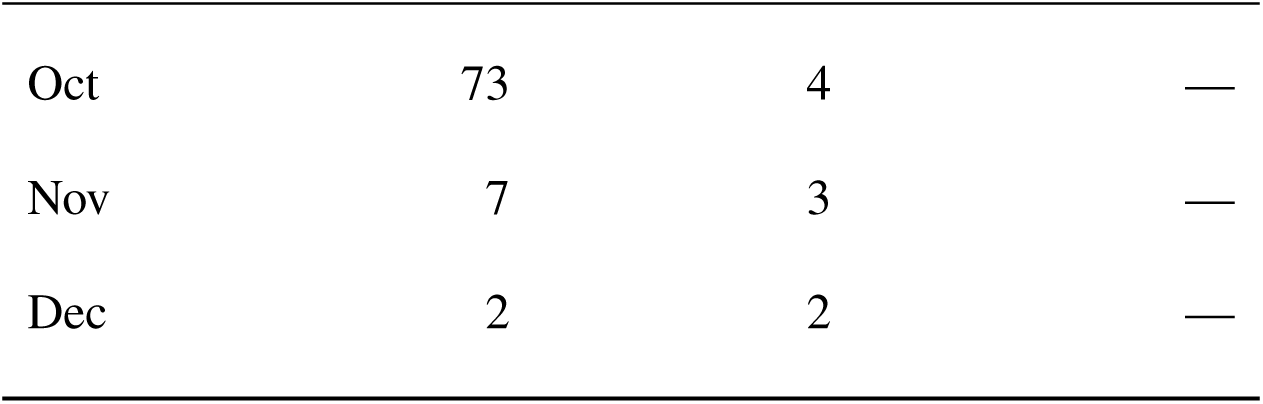
Monthly simulated whale-watching (WW) effort and detections, and survey detections.

| Month | WW presences | WW days out | Survey detections |
| --- | --- | --- | --- |
| Jan | 1 | 1 | — |
| Feb | 2 | 2 | — |
| Mar | 8 | 3 | — |
| Apr | 9 | 4 | — |
| May | 16 | 5 | — |
| Jun | 44 | 6 | — |
| Jul | 151 | 7 | — |
| Aug | 163 | 6 | 79 |
| Sep | 147 | 5 | — |
| Oct | 73 | 4 | — |
| Nov | 7 | 3 | — |
| Dec | 2 | 2 | — |

**Table S4:**
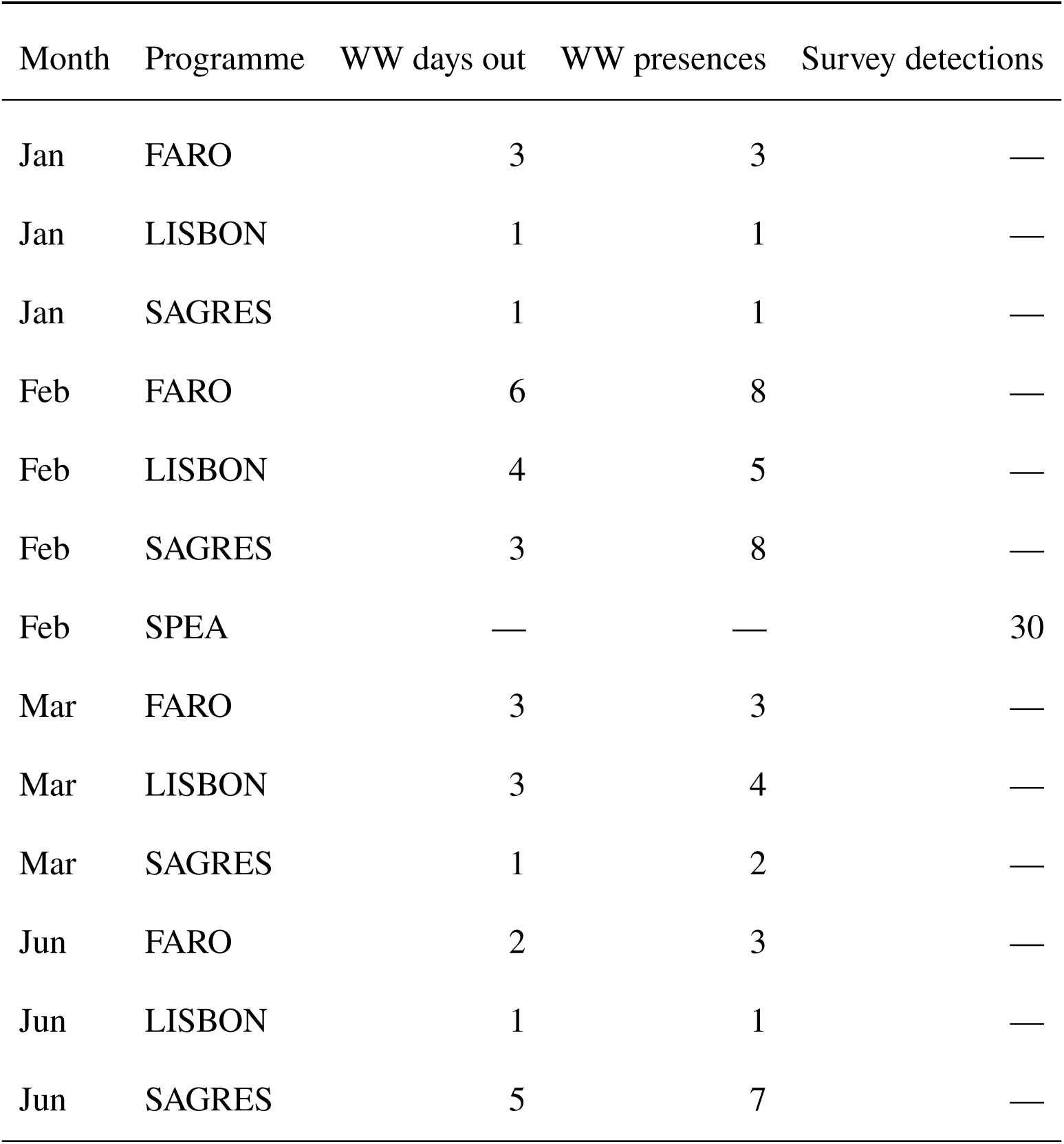
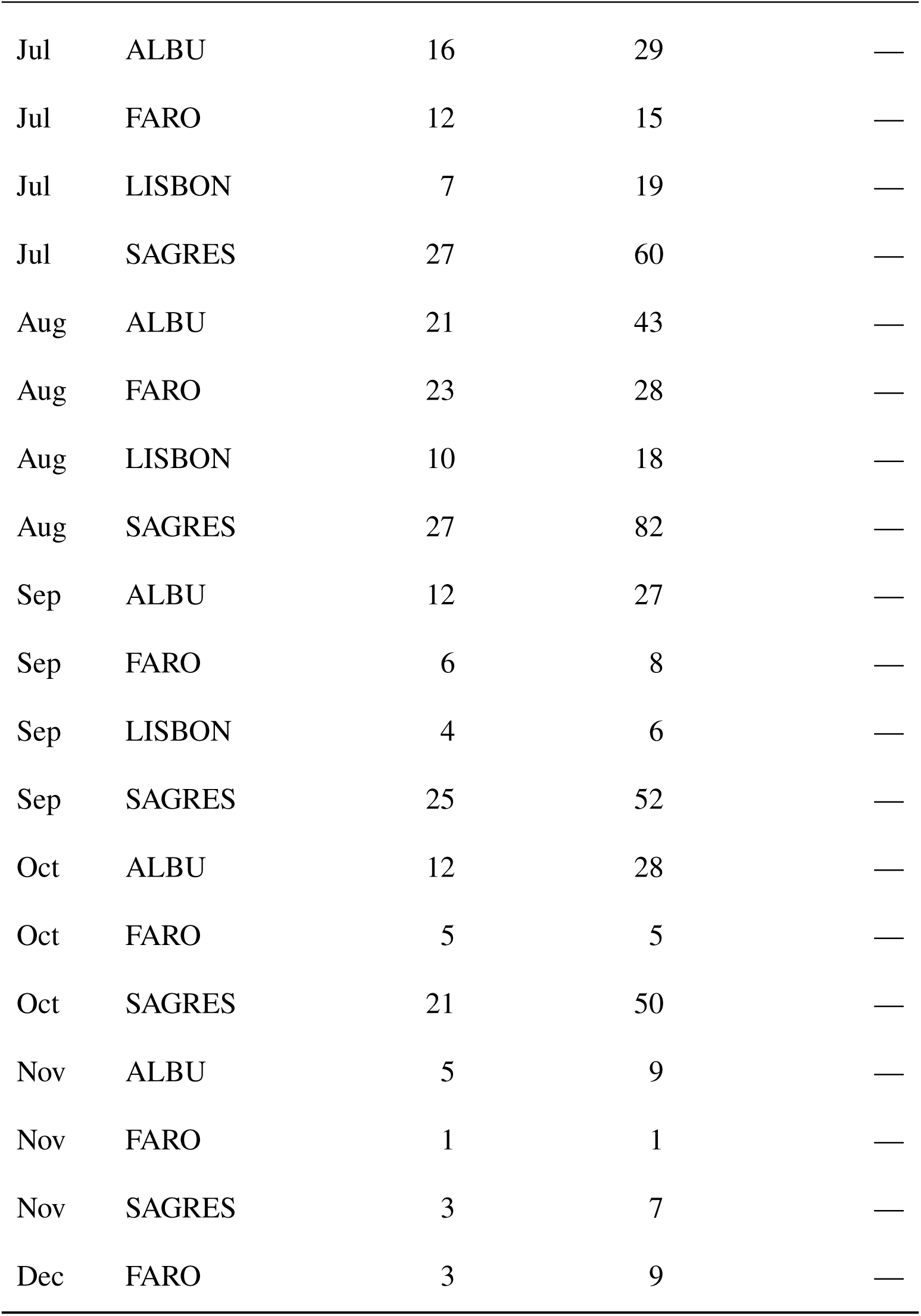
Monthly whale-watching (WW) effort and detections per operator, and survey detections (SPEA) for the case study.

| Month | Programme | WW days out | WW presences | Survey detections |
| --- | --- | --- | --- | --- |
| Jan | FARO | 3 | 3 | — |
| Jan | LISBON | 1 | 1 | — |
| Jan | SAGRES | 1 | 1 | — |
| Feb | FARO | 6 | 8 | — |
| Feb | LISBON | 4 | 5 | — |
| Feb | SAGRES | 3 | 8 | — |
| Feb | SPEA | — | — | 30 |
| Mar | FARO | 3 | 3 | — |
| Mar | LISBON | 3 | 4 | — |
| Mar | SAGRES | 1 | 2 | — |
| Jun | FARO | 2 | 3 | — |
| Jun | LISBON | 1 | 1 | — |
| Jun | SAGRES | 5 | 7 | — |
| Jul | ALBU | 16 | 29 | — |
| Jul | FARO | 12 | 15 | — |
| Jul | LISBON | 7 | 19 | — |
| Jul | SAGRES | 27 | 60 | — |
| Aug | ALBU | 21 | 43 | — |
| Aug | FARO | 23 | 28 | — |
| Aug | LISBON | 10 | 18 | — |
| Aug | SAGRES | 27 | 82 | — |
| Sep | ALBU | 12 | 27 | — |
| Sep | FARO | 6 | 8 | — |
| Sep | LISBON | 4 | 6 | — |
| Sep | SAGRES | 25 | 52 | — |
| Oct | ALBU | 12 | 28 | — |
| Oct | FARO | 5 | 5 | — |
| Oct | SAGRES | 21 | 50 | — |
| Nov | ALBU | 5 | 9 | — |
| Nov | FARO | 1 | 1 | — |
| Nov | SAGRES | 3 | 7 | — |
| Dec | FARO | 3 | 9 | — |

**Figure S7.**
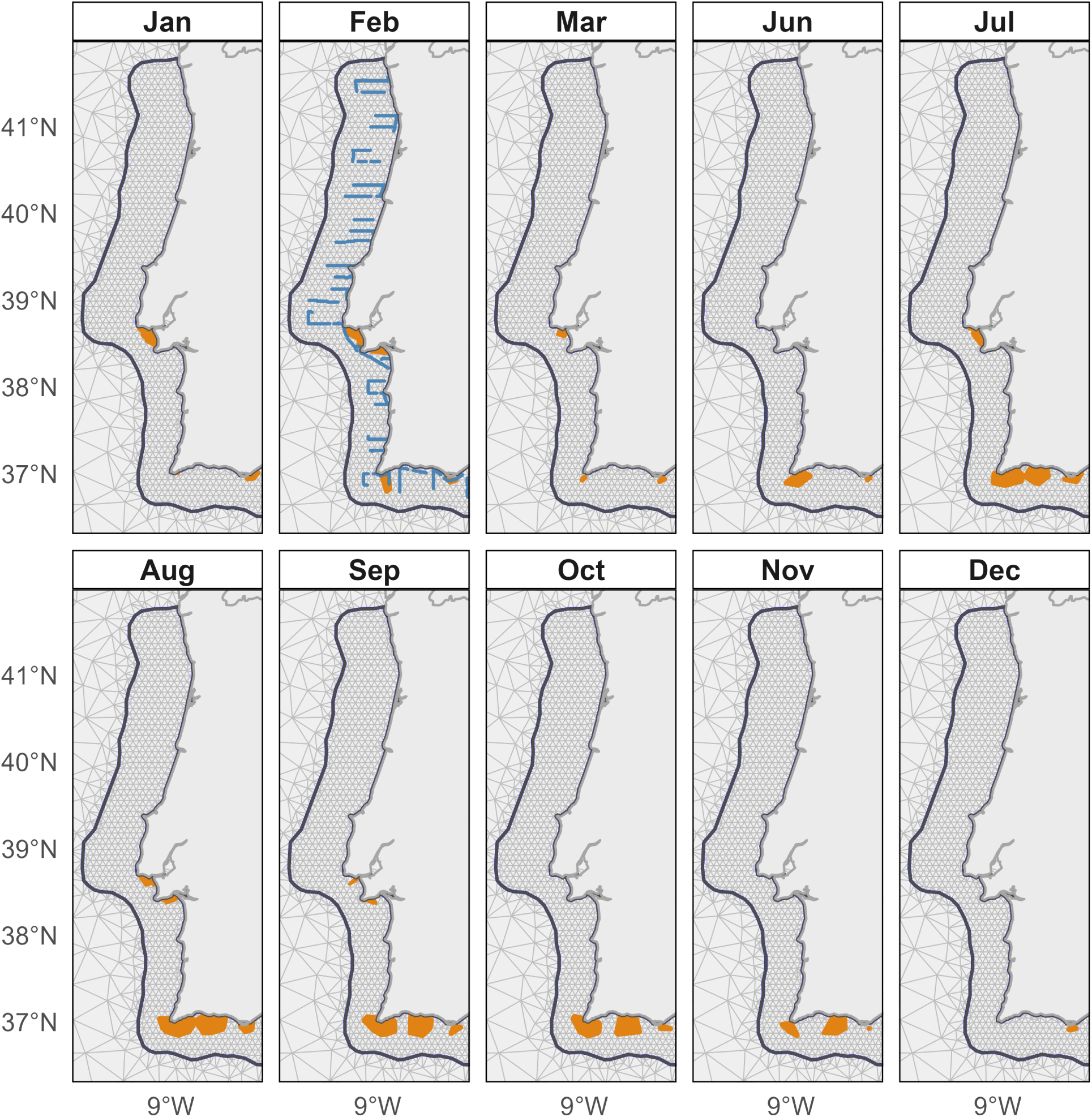
Monthly sampling effort for the common dolphin case study (2020). The grey triangulated background is the SPDE mesh (outer mesh cropped to keep effort visible). Survey effort from the SPEA line transect (February only) is shown in blue. WW effort is shown in orange. Only months with data are displayed.

**Figure S8.**
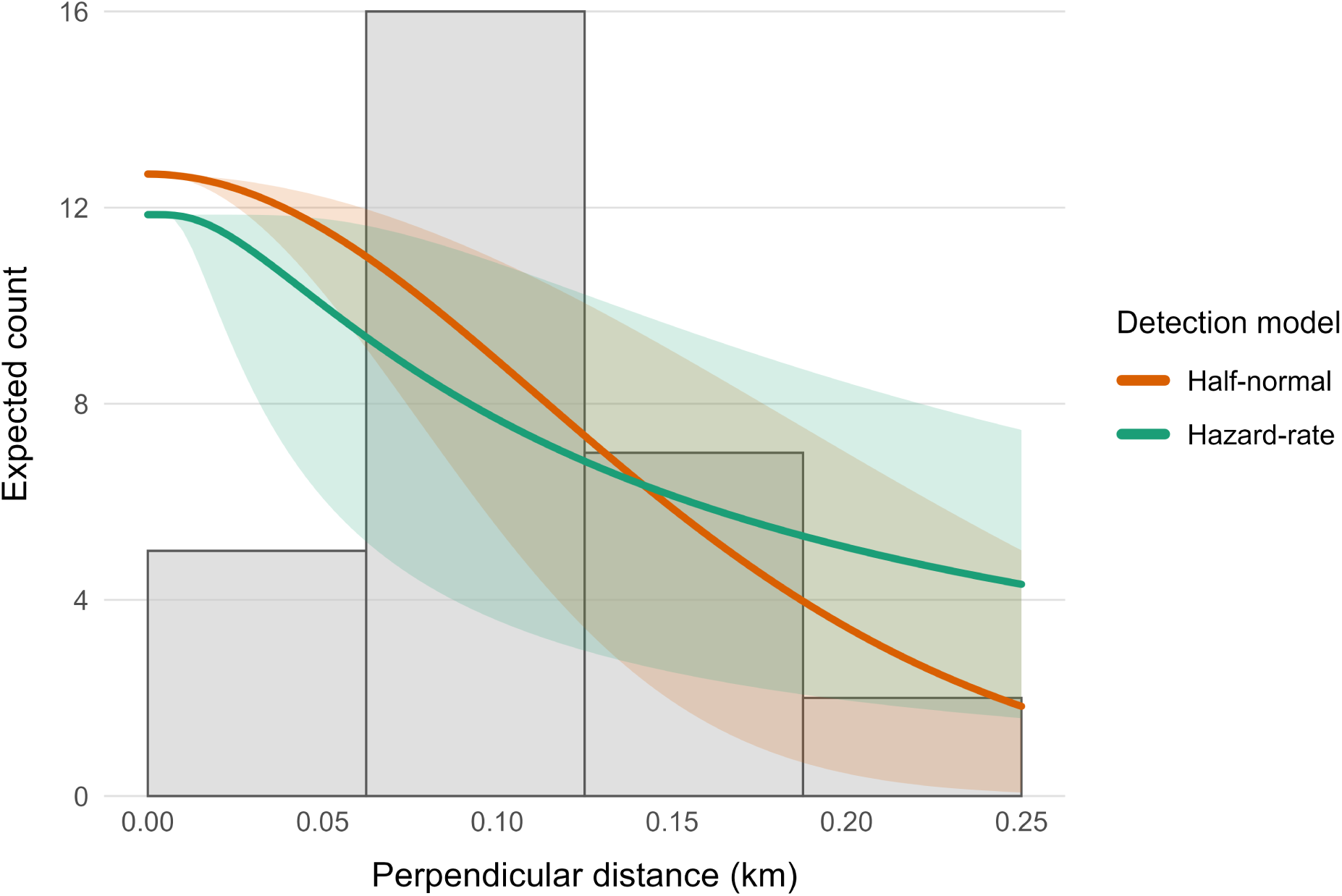
Detection model comparison for the survey data. Histogram of observed perpendicular distances (0–0.25 km; 4 bins) overlaid with expected bin counts from the fitted half-normal (orange) and hazard-rate (green) models; shaded ribbons show 95% credible bands. Despite the apparent misfit in the first bin, we opted against left truncation given the scope of this paper.

**Figure S9.**
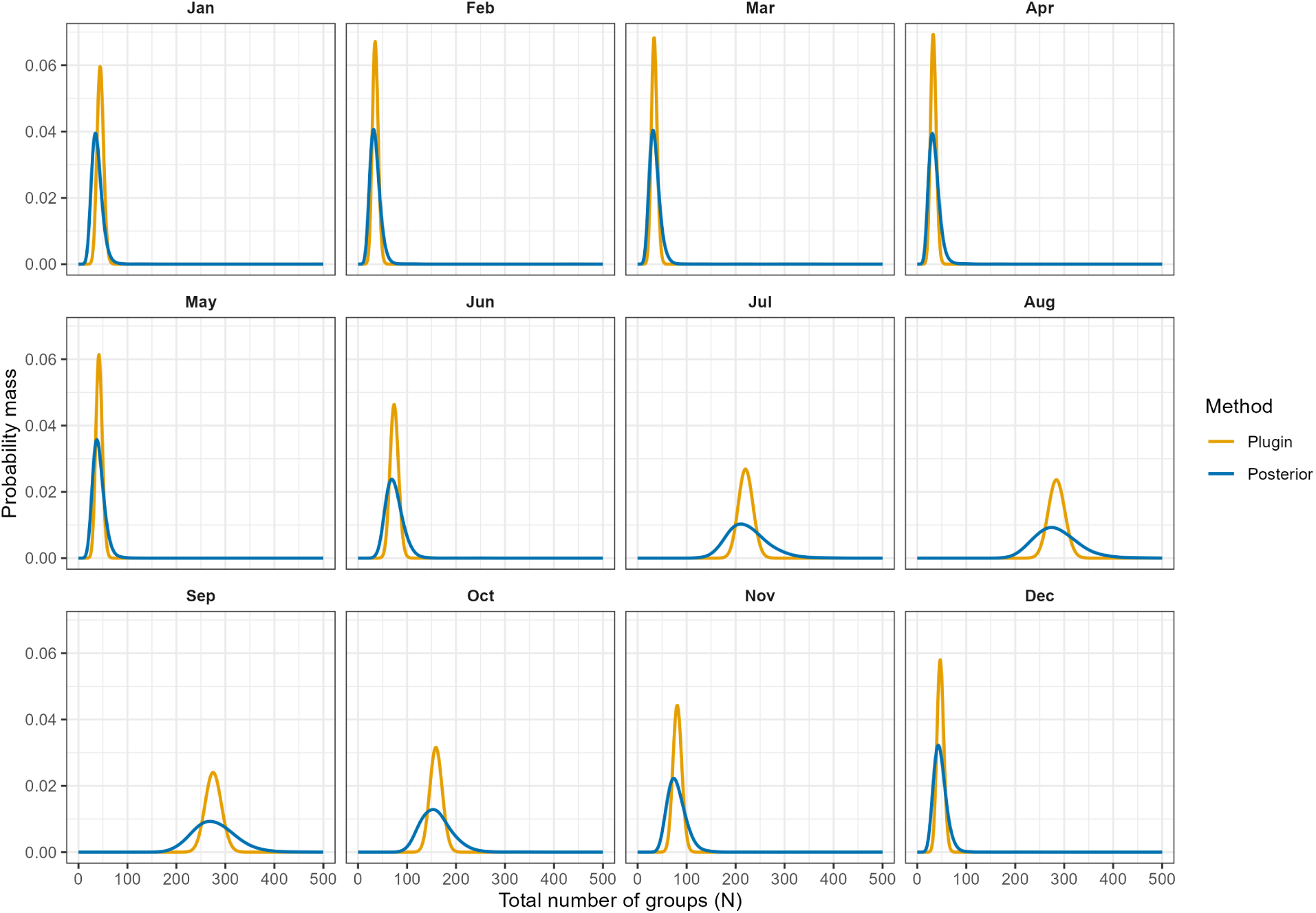
Monthly posterior predictive distributions compared with a plug-in Poisson approximation that conditions on the posterior mean expected abundance.

**Figure S10.**
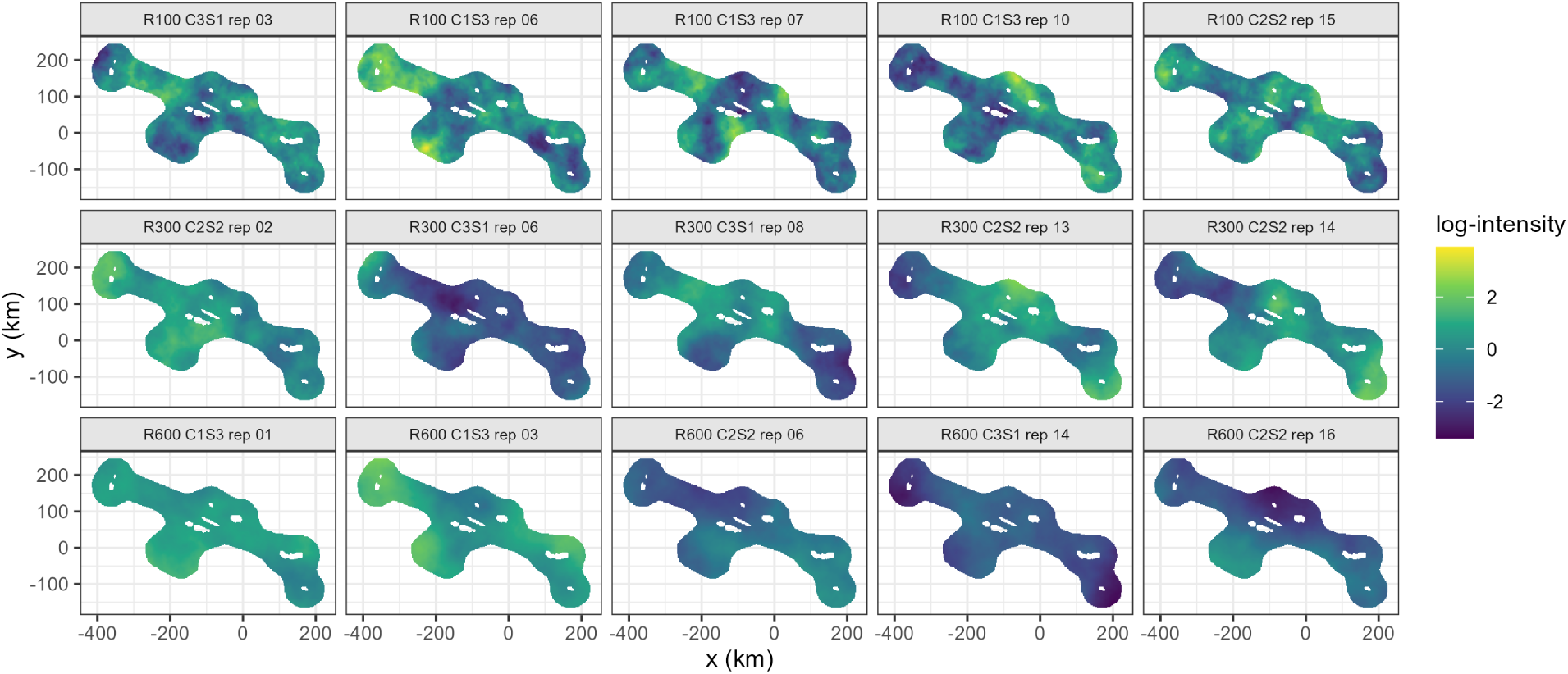
Simulated Gaussian random fields (GRFs) used as residual structure in the multi-scenario simulation. Each panel shows one realisation of the barrier-aware Matérn SPDE field; rows vary the water correlation range (100, 300, 600 km) and columns show five randomly selected replicates per range.

**Figure S11.**
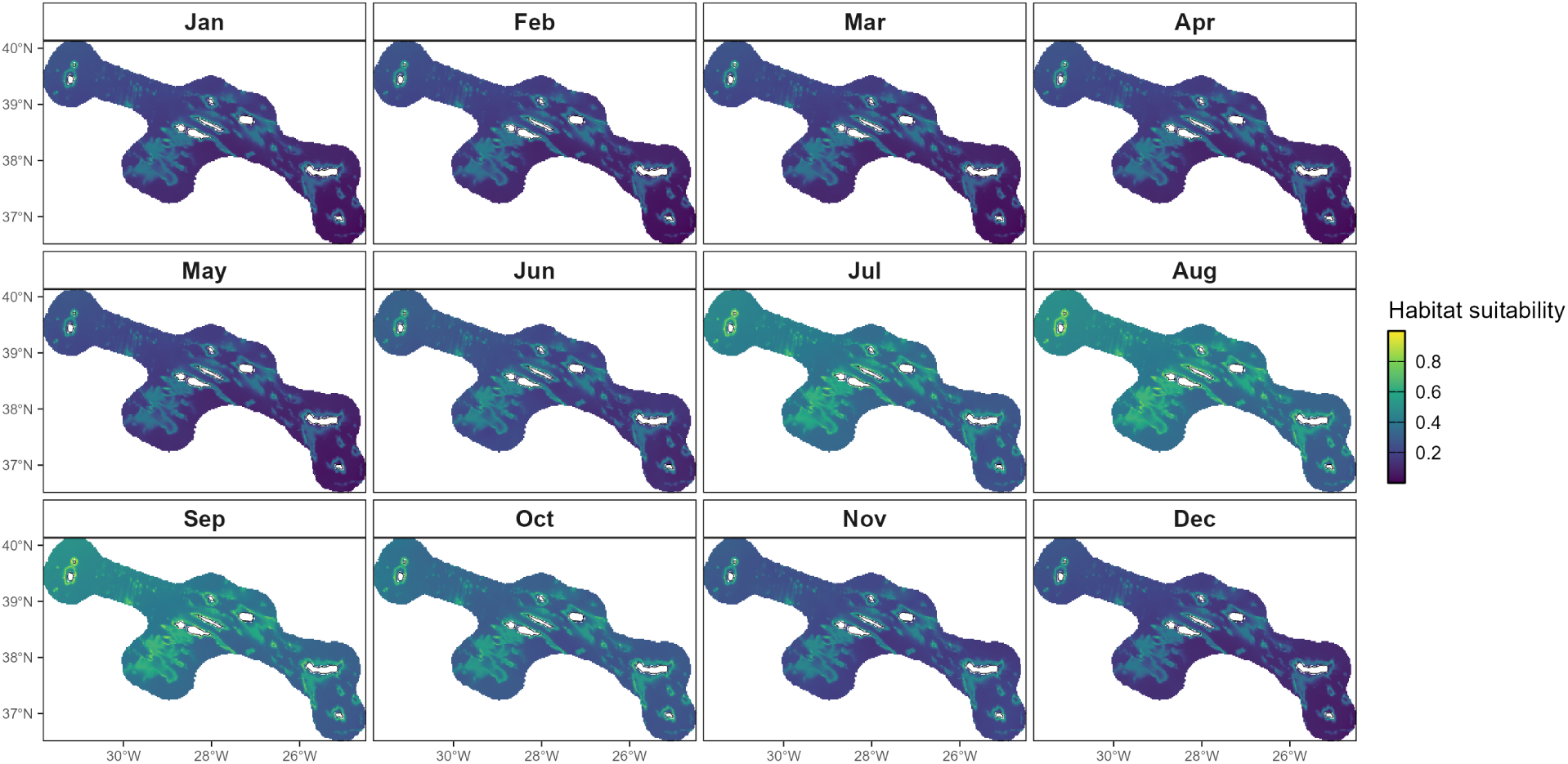
Monthly habitat suitability (0–1) for the Azores, used as the point-generating surface in the simulation.

**Figure S12.**
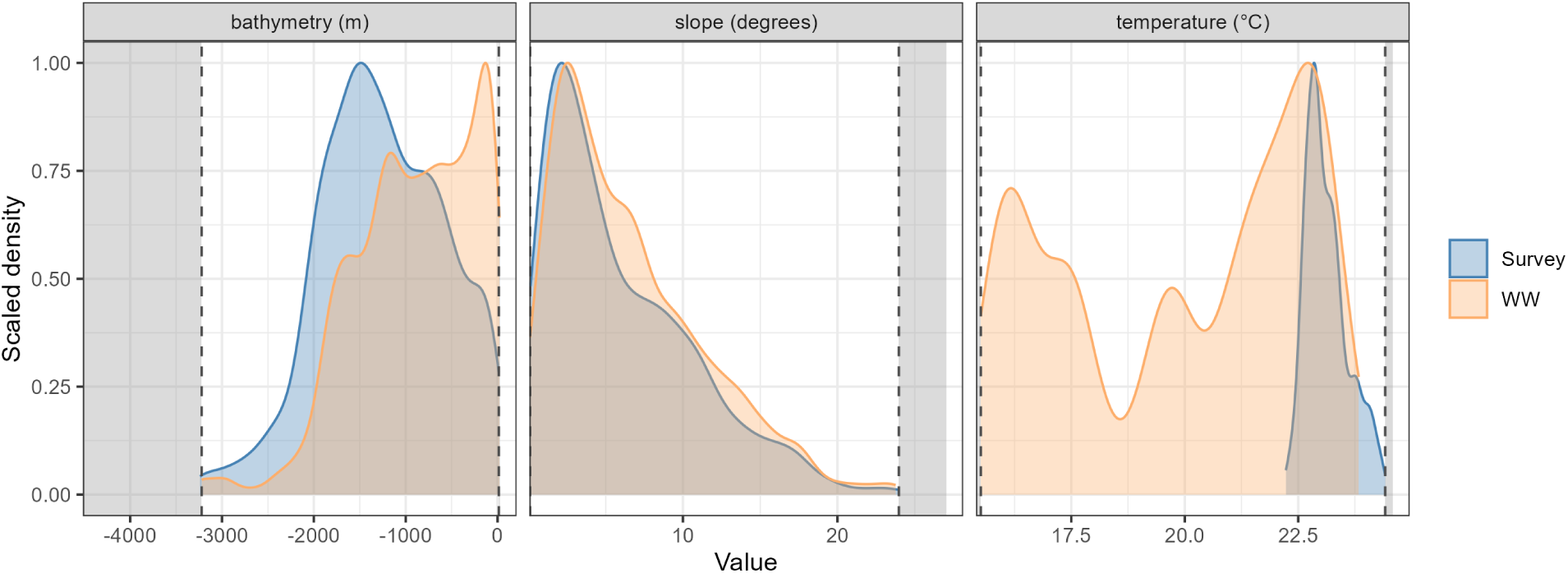
Effort coverage in covariate space for the simulation. Kernel densities of covariate values at integration points for the survey (blue; August line-transect) and whale-watching effort (orange; pooled across months). Panels show bathymetry, slope, and temperature. Dashed vertical lines mark the min–max range covered by the survey and whale-watching effort. Grey shading indicates regions of the prediction domain outside the covariate coverage where the model relies on extrapolation.

**Figure S13.**
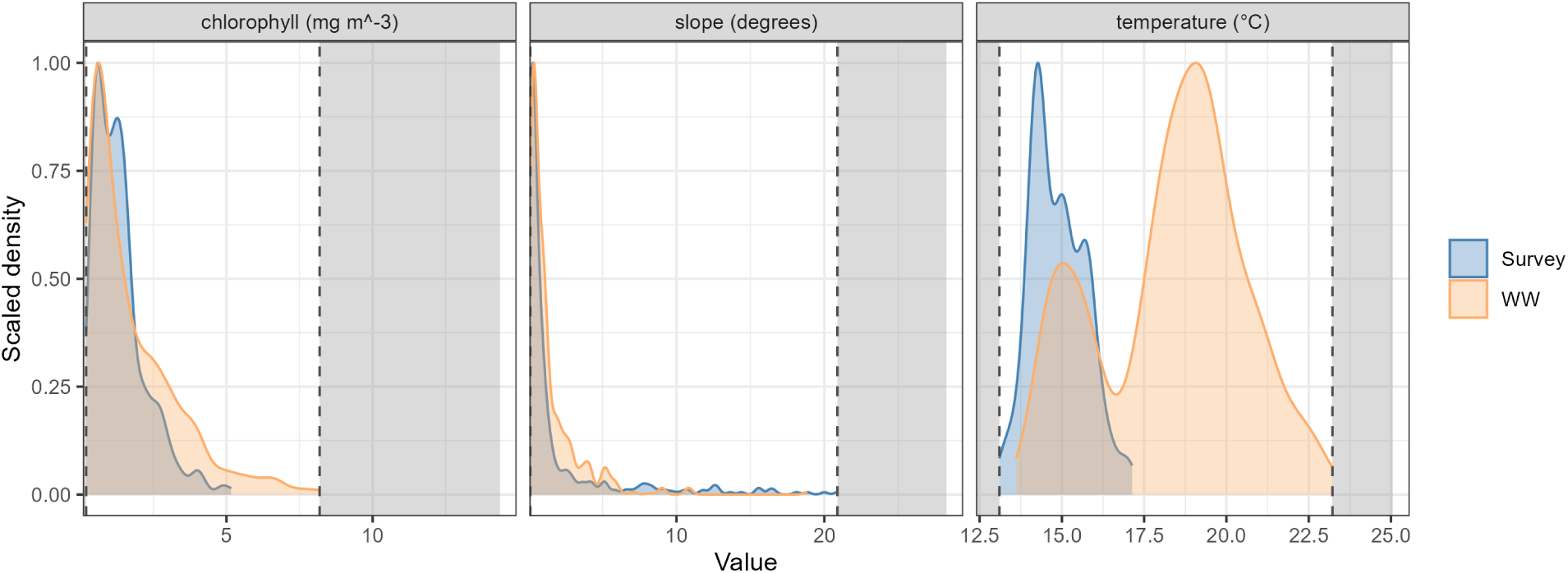
Effort coverage in covariate space for the case study. Kernel densities of covariate values at integration points for the survey (blue; February line-transect) and pooled whale-watching programmes (orange). Panels show slope, temperature, and chlorophyll. Dashed vertical lines mark the min–max range covered by the survey and whale-watching effort. Grey shading indicates regions of the prediction domain outside the covariate coverage where the model relies on extrapolation.

**Figure S14.**
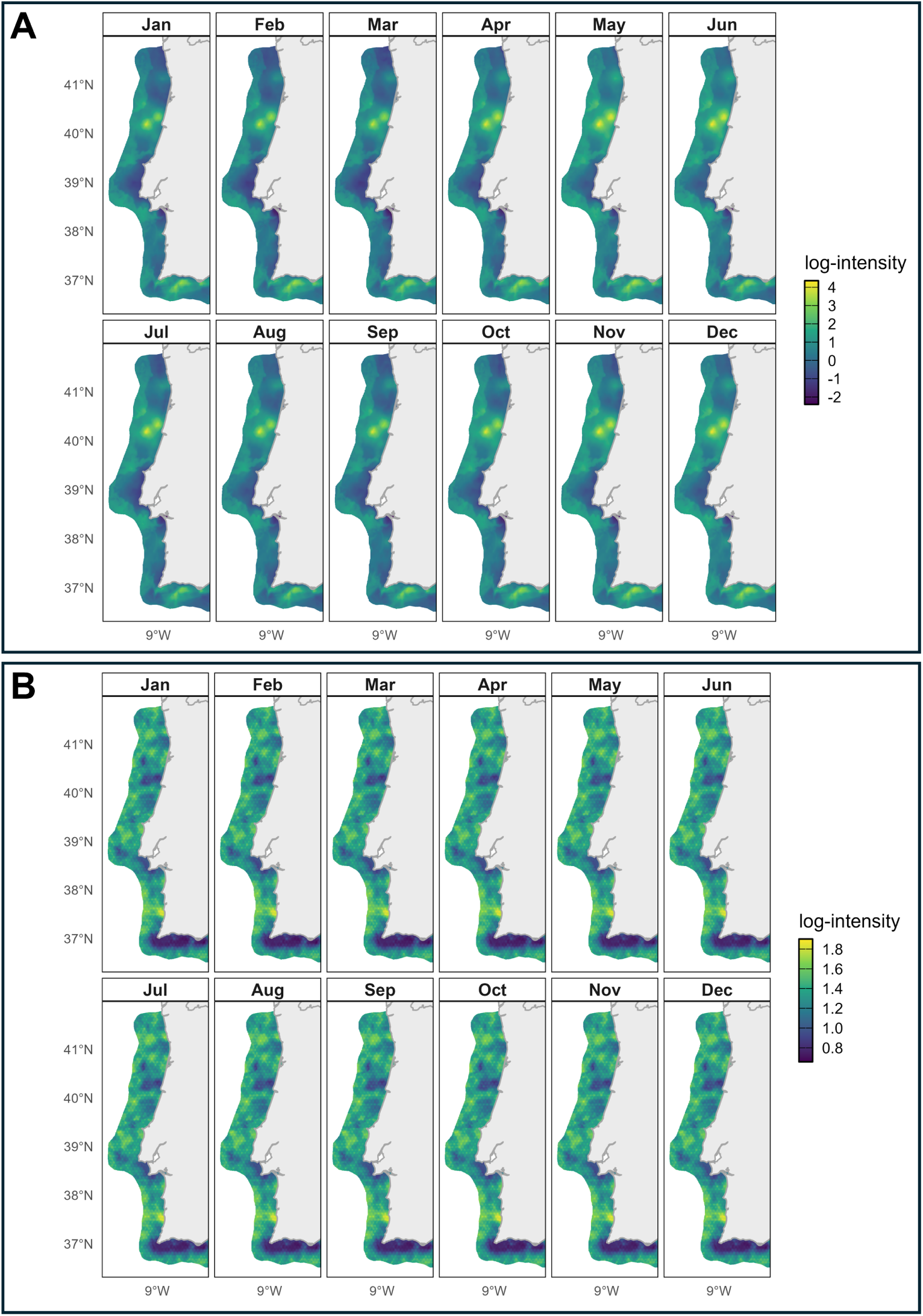
Monthly dynamics of the linear predictor for the Portugal case study. **(A)** Posterior mean of log-intensity by month. **(B)** Posterior standard deviation by month.

## References

Bachl, F. E., F. Lindgren, D. L. Borchers, and J. B. Illian. 2019. “inlabru: an R Package for Bayesian Spatial Modelling from Ecological Survey Data.” Methods in Ecology and Evolution 10 (6): 760–766.

Bakka, H., J. Vanhatalo, J. B. Illian, D. Simpson, and H. Rue. 2019. “Non-Stationary Gaussian Models with Physical Barriers.” Spatial Statistics 29:268–288.

Buckland, S. T., E. A. Rexstad, T. A. Marques, and C. S. Oedekoven. 2015. Distance Sampling: Methods and Applications. Methods in Statistical Ecology. Cham: Springer International Publishing.

Buckland, S. T., D. R. Anderson, K. P. Burnham, J. L. Laake, D. L. Borchers, and L. Thomas. 2001. Introduction to Distance Sampling: Estimating Abundance of Biological Populations. Oxford: Oxford University Press.

Diggle, P. J., R. Menezes, and T.-L. Su. 2010. “Geostatistical Inference under Preferential Sampling.” Journal of the Royal Statistical Society: Series C (Applied Statistics) 59 (2): 191–232.

Elith, J., and J. R. Leathwick. 2009. “Species Distribution Models: Ecological Explanation and Prediction Across Space and Time.” Annual Review of Ecology, Evolution, and Systematics 40:677–697.

Farr, M. T., D. S. Green, K. E. Holekamp, and E. F. Zipkin. 2021. “Integrating Distance Sampling and Presence-Only Data to Estimate Species Abundance.” Ecology 102 (1): e03204.

Farr, M. T., E. R. Zylstra, L. Ries, and E. F. Zipkin. 2024. “Overcoming Data Gaps Using Integrated Models to Estimate Migratory Species’ Dynamics During Cryptic Periods of the Annual Cycle.” Methods in Ecology and Evolution 15 (2): 413–426.

Fretwell, P. T., I. J. Staniland, and J. Forcada. 2014. “Whales from Space: Counting Southern Right Whales by Satellite.” PLoS One 9:e88655.

Gilbert, N. A., B. R. Amaral, O. M. Smith, P. J. Williams, S. Ceyzyk, S. Ayebare, K. L. Davis, W. Leuenberger, J. W. Doser, and E. F. Zipkin. 2024. “A century of statistical Ecology.” Ecology 105 (6): e4283.

Hammond, P. S., P. Berggren, H. Benke, D. Borchers, A. Collet, M. Heide-Jørgensen, S. Heimlich, A. Hiby, M. Leopold, and N. Øien. 2002. “Abundance of Harbour Porpoise and Other Cetaceans in the North Sea and Adjacent Waters.” Journal of Applied Ecology 39 (2): 361–376.

Hammond, P. S., K. Macleod, P. Berggren, D. L. Borchers, L. Burt, A. Cañadas, G. Desportes, et al. 2013. “Cetacean abundance and distribution in European Atlantic shelf waters to inform conservation and management.” Biological Conservation 164:107–122.

Hazen, E. L., K. L. Scales, S. M. Maxwell, D. K. Briscoe, H. Welch, S. J. Bograd, H. Bailey, et al. 2018. “A Dynamic Ocean Management Tool to Reduce Bycatch and Support Sustainable Fisheries.” Science Advances 4 (5): eaar3001.

Illian, J. B., A. Penttinen, H. Stoyan, and D. Stoyan. 2008. Statistical Analysis and Modelling of Spatial Point Patterns. Chichester: John Wiley & Sons, 2008.

Kaschner, K., N. J. Quick, R. Jewell, R. Williams, and C. M. Harris. 2012. “Global Coverage of Cetacean Line-Transect Surveys: Status Quo, Data Gaps and Future Challenges.” PLOS ONE 7, no. 9 (2012): e44075.

Kéry, M., J. A. Royle, T. Hallman, W. D. Robinson, N. Strebel, and K. F. Kellner. 2024. “Integrated Distance Sampling Models for Simple Point Counts.” Ecology 105, no. 5 (2024): e4292.

Klaassen, M., T. A. Marques, F. Alves, and M. Fernandez. 2025. “Trends in Marine Species Distribution Models: a Review of Methodological Advances and Future Challenges.” Ecography, e07702.

Krainski, E. T., F. Lindgren, and H. Rue. 2025. INLAspacetime: Spatial and Spatio-Temporal Models using INLA. R package.

Lindgren, F. 2025. fmesher: Triangle Meshes and Related Geometry Tools. R package.

Lindgren, F., H. Rue, and J. Lindström. 2011. “An Explicit Link Between Gaussian Fields and Gaussian Markov Random Fields: the Stochastic Partial Differential Equation Approach.” Journal of the Royal Statistical Society: Series B (Statistical Methodology) 73 (4): 423–498.

Marques, T. A., L. Thomas, S. W. Martin, D. K. Mellinger, J. A. Ward, D. J. Moretti, D. Harris, and P. L. Tyack. 2013. “Estimating Animal Population Density Using Passive Acoustics.” Biological Reviews 88:287–309.

Martino, S., D. S. Pace, S. Moro, E. Casoli, D. Ventura, A. Frachea, M. Silvestri, et al. 2021. “Integration of presence-only data from several sources: a case study on dolphins’ spatial distribution.” Ecography 44 (10): 1533–1543.

Miller, D. A. W., K. Pacifici, J. S. Sanderlin, and B. J. Reich. 2019. “The Recent Past and Promising Future for Data Integration Methods to Estimate Species’ Distributions.” Methods in Ecology and Evolution 10 (1): 22–37.

Pacifici, K., B. J. Reich, D. A. W. Miller, B. Gardner, G. Stauffer, S. Singh, A. McKerrow, and J. A. Collazo. 2017. “Integrating multiple data sources in species distribution modeling: a framework for data fusion.” Ecology 98 (3): 840–850.

Renner, I. W., and D. I. Warton. 2013. “Equivalence of MAXENT and Poisson Point Process Models for Species Distribution Modeling in Ecology.” Biometrics 69 (1): 274–281.

Rue, H., S. Martino, and N. Chopin. 2009. “Approximate Bayesian Inference for Latent Gaussian Models by Using Integrated Nested Laplace Approximations.” Journal of the Royal Statistical Society: Series B (Statistical Methodology) 71 (2): 319–392.

Seaton, F. M., S. G. Jarvis, and P. A. Henrys. 2024. “Spatio-Temporal Data Integration for Species Distribution Modelling in R-INLA.” Methods in Ecology and Evolution 15 (7): 1221–1232.

Simpson, D., H. Rue, A. Riebler, T. G. Martins, and S. H. Sørbye. 2017. “Penalising Model Component Complexity: A Principled, Practical Approach to Constructing Priors.” Statistical Science 32 (1): 1–28.

Alves, F., A. Alessandrini, A. Servidio, A. S. Mendonça, K. L. Hartman, R. Prieto, S. Berrow, et al. 2019. “Complex Biogeographical Patterns support an Ecological Connectivity Network of a Large Marine Predator in the North-East Atlantic.” Diversity and Distributions 25 (2): 269–284.

Bravington, M. V., D. L. Miller, and S. L. Hedley. 2021. “Variance Propagation for Density Surface Models.” *Journal of Agricultural*, Biological and Environmental Statistics 26 (2): 306–323.

Faustino, C. E. S., M. A. Silva, T. A. Marques, and L. Thomas. 2010. “Designing a Shipboard Line Transect Survey to Estimate Cetacean Abundance off the Azores Archipelago.” Arquipelago. Life and Marine Sciences 27:49–58.

Fernandez, M., N. Sillero, and C. Yesson. 2022. “To Be or Not to Be: the Role of Absences in Niche Modelling for Highly Mobile Species in Dynamic Marine Environments.” Ecological Modelling 471:110040.

GBIF. 2025. “Global Biodiversity Information Facility.” Accessed 15 Feb 2025. https://www.gbif.org.

González García, L., M. Fernandez, and J. M. N. Azevedo. 2023. “MONICET: The Azores Whale Watching Contribution to Cetacean Monitoring.” Biodiversity Data Journal 11:e106991.

Leroy, B., C. N. Meynard, C. Bellard, and F. Courchamp. 2016. “virtualspecies, an R Package to Generate Virtual Species Distributions.” Ecography 39 (6): 599–607.

